# A supragranular nexus for the effects of neocortical beta events on human tactile perception

**DOI:** 10.1101/750992

**Authors:** Robert G. Law, Sarah Pugliese, Hyeyoung Shin, Danielle Sliva, Shane Lee, Samuel Neymotin, Christopher Moore, Stephanie R. Jones

**Affiliations:** Brown University Department of Neuroscience; Providence RI; Center for Neurorestoration and Neurotechnology; Providence VA Medical Center, Providence RI; Stanley Center for Psychiatric Research; The Broad Institute of MIT and Harvard, Cambridge MA; Nathan S. Kline Institute for Psychiatric Research; Orangeburg NY

## Abstract

Transient neocortical events with high spectral power in the 15–29Hz beta band are among the most reliable predictors of sensory perception: High prestimulus beta event rates in primary somatosensory lead to sensory suppression, most effective at 100–300ms prestimulus latency. However, the synaptic and neuronal mechanisms inducing beta’s perceptual effects have not been completely localized. We combined human MEG with neural modeling designed to account for these macroscale signals to interpret the cellular and circuit mechanisms that underlie the influence of beta on tactile detection. Extending prior studies, we modeled the hypothesis that higher-order thalamic bursts, sufficient for beta event generation in cortex, recruit supragranular GABA_B_ inhibition acting on a 300ms time scale to suppress sensory information. Consistency between model and MEG data supported this hypothesis and led to a further prediction, validated in our data, that stimuli are perceived when beta events occur simultaneously with tactile stimulation. The post-event suppressive mechanism explains an array of studies that associate beta with decreased processing, while the during-event mechanism may demand a reinterpretation of the role of beta events in the context of coincident timing.

**Significance statement:** Somatosensory beta events – transient 15-29Hz oscillations in electromagnetic recordings – are thought to be generated when “top-down” bursts of spikes presumably originating in higher-order thalamus arrive in upper layers of somatosensory cortex. Physiological evidence had shown that the immediate action of these top-down projections should be excitatory; however, after a beta event, sensory perception is noticeably inhibited for approximately 300ms. The source of this post-event sensory suppression, in particular, had been unresolved. Using a detailed computational model of somatosensory cortex, we find evidence for the hypothesis that these bursts couple indirectly to GABA_B_ inhibition in upper layers of cortex, and that beta events first briefly disinhibit sensory relay before a longer period of inhibition.

## Introduction

Brain rhythms need not be recurrent oscillations, raising subtle questions as to precisely if and how a transient, even monocyclic “rhythm” can causally influence sensory and cognitive processing (1–3). The fact that brain dynamics are nonstationary (4) — with time-localized spectral energy on individual trials non-uniformly contributing to time-averaged spectral power — has become a focus of a number of research efforts (1–3, 5–8). Although transient rhythms have long been a central focus in sleep research (9), this more recent line of questioning concerns awake, behaving animals. Here, modern consensus holds that many brain rhythms tend to bubble in and out of existence, appearing as sparse, energetically concentrated *events* (cf. *bursts*^*^) that can have marked influence on waking perception and behavior.

Transient beta band (15–29Hz) rhythms are one of the most well-established *event-like* rhythms, observed in sensory (5, 10), motor (11, 12), prefrontal (6, 10, 13), and subcortical regions (7), and across species and recording modalities (5). Modulation of beta events, as quantified in event rate or time-dependent probability density, are associated with memory processes (6, 13), sensory perception (5) and motor action in both health and disease (12, 14–16). Yet, the neural mechanisms by which beta events influence behavior are currently unknown, and the cognitive and perceptual roles of beta remain in debate (17–19).

We had previously observed that the rate and timing of beta events in magnetoencephalographic (MEG) and local field potential (LFP) recordings from primary somatosensory cortex (SI) predict threshold-level tactile perception (5, 10). Two or more events in a one-second prestimulus period decreased perception, and, despite a typical event lifetime of less than 150ms, perception was strongly impaired 100–300ms following a beta event (5).

Our first goal was therefore to understand beta’s role in this perceptual suppression, and we extended neural modeling designed to interpret the circuit origin of human MEG/EEG signals based on their best-known electromagnetic sources: intracellular current flows driven by active membranes in pyramidal neurons (10, 20–23). Here, we use the model to identify cellular and circuit dynamics by which human SI beta events may influence threshold-level perception — and thereby affect tactile evoked responses (20, 21, 24) — by modeling both endogenous beta events and subsequent evoked responses. We first refined our model of the generation of beta events (10), where bursts of excitatory drive to supragranular layers, most likely originating in higher-order thalamus, are *sufficient* to generate strong tuft excitation. This excitation leads to downward currents *necessary* for beta event generation in humans.

Based on observation that inhibitory effects of beta events last ~100-300ms, we hypothesized that slow inhibitory currents at the GABA_B1a_ timescale (25) are also recruited by these distal-targeting bursts. This hypothesis is consistent with the locus of GABA_B1a_-targeting neurogliaform cells in supragranular cortical layers that can somatically inhibit SI pyramidal neurons in L2/3 (26, 27), and complements prior studies showing a direct influence of supragranular GABA_B_ inhibition on tactile detection (28). By extending our model to incorporate supragranular GABA_B_-mediated inhibition, we were able to account for beta’s suppressive effects on tactile evoked responses and perception in agreement with our MEG data.

With a model of post-event effects in hand, we then considered what occurs when a stimulus arrives *during* a beta event. Our modeling revealed that the distal drive that generates beta events causes sufficient excitation to yield a brief time window where beta events cause a bias toward detection before long-timescale inhibition is engaged. We find evidence for this rare effect, which depends on consideration of event sparsity and narrowband beta phase.

Overall, the consistency between model results and human recordings indicates that beta events in SI can briefly act to promote stimulus information before a longer period of GABA_B_ inhibition that dominates the average behavioral effect. These results are discussed in relation to an array of studies that associate beta with decreased processing, as well as literature that suggest beta may reflect timed prediction and learning of cognitively relevant stimuli.

## Results

### A Single Narrowband Prestimulus Beta Event Decreases Tactile Detection

We have previously shown that two or more 15–29Hz SI beta events yield a strong bias toward non-detection of a perceptual threshold-level tactile stimulus if they occur in the 1000ms pre-stimulus time window — a consistent finding across mouse LFP and human MEG (5). These events were defined as periods with power above a 6x-median threshold in the time-frequency plane (see Experimental Procedures). Though the presence of a single 15–29Hz event did not significantly influence detection (5), we observed that human beta events tend to concentrate in a narrowband near 20Hz (e.g. Figure 1A); when peak power was found in the 20-22Hz range, a single prestimulus beta event was enough to impair tactile detection significantly (Figure 1B). We therefore partitioned trials into those with at least one 20–22Hz event (*event* trials) and those without (*no-event* trials). A representative beta event is shown (Figure 1A), with an individual beta cycle highlighted for comparison to the beta event model to follow. Event trials had reduced detection probability across subjects (8/10 participants; p < 0.01; Wilcoxon rank-sum test; in total: 502 event trials (295 miss, 207 hit) and 1498 no-event trials (705 miss, 793 hit)), with individual changes in detection probability shown in Figure 1B (median reduction 10%; maximum reduction 28%).

**Figure 1:**
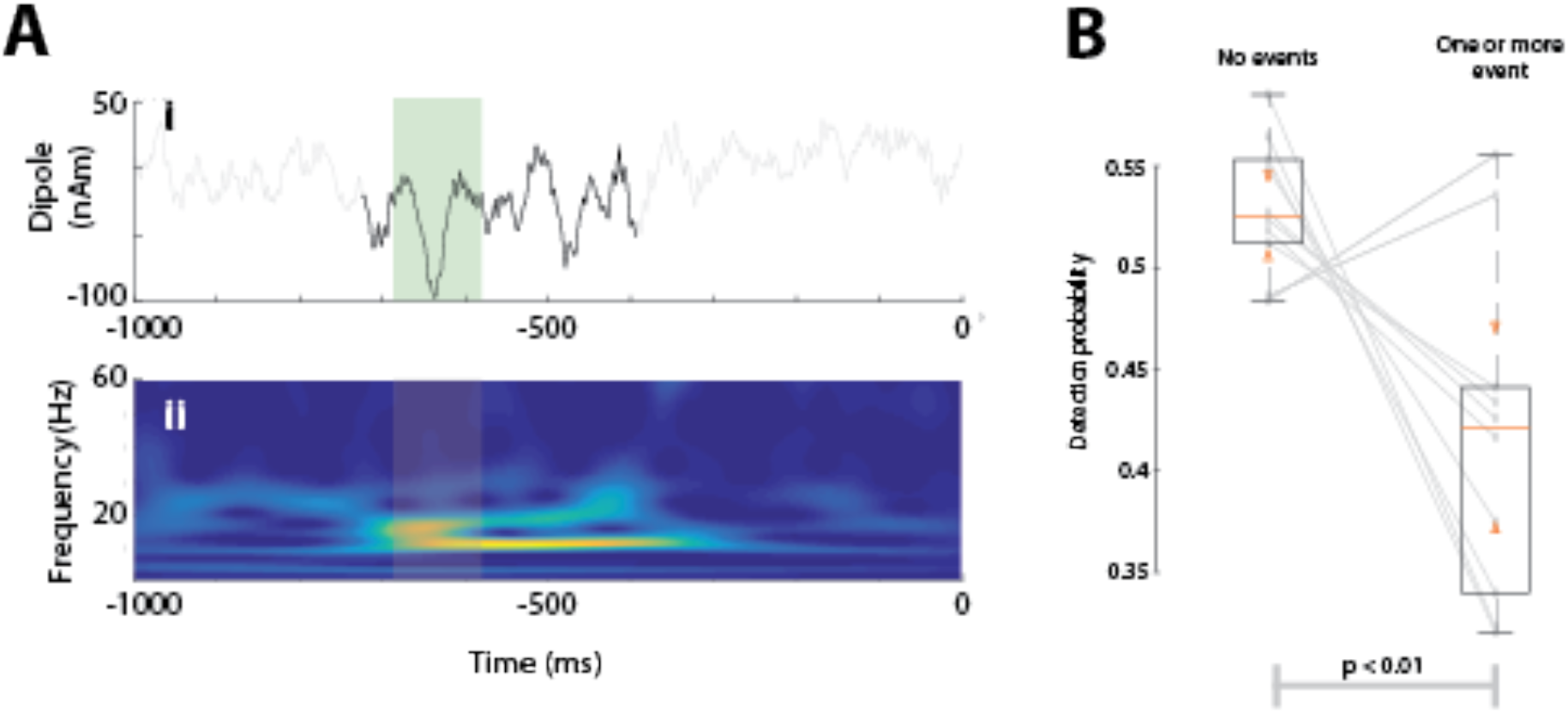
Prestimulus narrowband-powered (20-22Hz) beta events are associated with lower near-threshold detection probabilities in human SI. **A)** Example trial with a 20-22Hz beta event represented in **i)** the time-domain with beta event in black (model beta cycle is highlighted in blue) and **ii)** as a wavelet spectrogram with event timespan highlighted. **B)** Detection probabilities for all participants, with boxes showing interquartile ranges. Orange bars and triangles represent median and comparison intervals respectively; see Methods. A 20-22Hz beta event occurring in the prestimulus period reduces detection probability (p < 0.01; Wilcoxon rank-sum test) by approximately 10% on average, and on an individual basis a prestimulus beta event alters detection probability in a range of −28% to +7%. The two outliers for whom beta events increased detection probability also had event-timing distributions differing from the remainder of the group (data not shown).

### 80-110ms is a critical time period for beta event effects on detection

We then set out to understand if and how SI beta events influence tactile detection. If a causal relationship were to exist, we would expect prestimulus beta events to directly alter the poststimulus tactile evoked response. Furthermore, we might expect such time-localized differences between event and no-event trials to match differences between hit and miss trials.

We first compared early SI tactile-evoked responses (0-140ms) between event and no-event trials (Figure 2A). We found two main effects of prestimulus beta events on evoked responses. Prestimulus events led to smaller amplitudes of evoked response components from ~98-113ms (Figure 2A; yellow bars; p < 0.05; permutation test, uncorrected) and a slower rise after the prominent negative peak (~70ms). Quantifying this with the time-derivative of the signals showed a difference just before the amplitude difference, around ~80-95ms post-stimulus, (Figure 2A inset).

**Figure 2:**
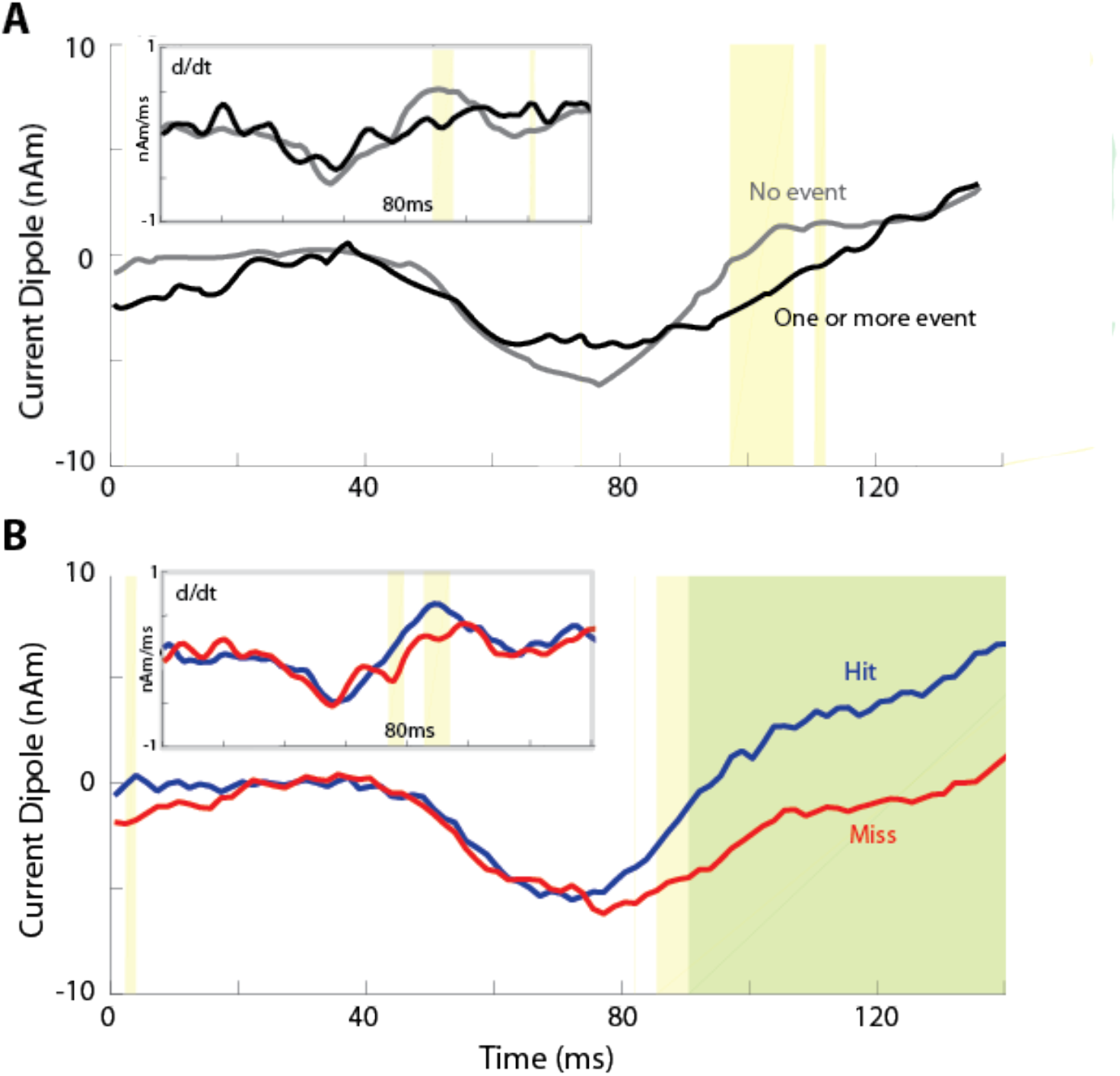
Human tactile evoked responses conditioned on prestimulus beta events and perception. **A)** Mean evoked responses for trials with or without a prestimulus beta event. **B)** Mean evoked responses for detected (“hit”) and nondetected (“miss”) trials. Significance is reported both pointwise in yellow (p < 0.05; permutation test) and after FWER-correction in green (α = 0.05; modified max-T test; see Supplementary Methods). *Insets:* Mean evoked response time-derivatives (after total-variation regularization; see Supplementary Methods).

We then compared evoked responses in hit and miss trials to determine if there were detection-related differences consistent with those occurring after beta events. The two beta event effects described above were also observed when partitioning by detection, consistent with our prior reports (21). Miss trials had smaller amplitudes from ~87-140ms (Figure 2B; yellow: p < 0.05; permutation test, uncorrected; green: alpha = 0.05; FWER-corrected permutation test), and the signal time-derivative showed a difference between hit and miss trials around ~75-97ms post-stimulus (Figure 2B; inset).

We conclude that tactile detection and SI beta events have intersecting intervals of post-stimulus effect, which provides support for a modeling hypothesis that beta events impact detection by affecting SI dynamics in the 80-110ms post-stimulus period.

### Resting model reproduces no-event trial averages

Having established a quantifiable association between beta events and non-detection, we next sought to understand circuit mechanisms by which this could occur — and why these mechanisms might correspond to impaired detection. To do so, we applied our biophysically principled model of a SI circuit that is specifically designed to simulate current dipole activity from the intracellular current flow in cortical pyramidal neurons, as in prior studies (20, 21, 23) (see Experimental Procedures, SI circuit (Figure 10)). We simulated a tactile evoked response without a preceding beta event, and during or after a simulated beta event, and examined the underlying circuit activity, as detailed below.

To reproduce the tactile-evoked response in our model, we simulated a sequence of external drives to the local network through layer-specific pathways based on known sensory evoked inputs to SI (Figure 3A - simulation with no beta event influence, Table 1), as in our prior studies (20, 21). Sensory input first arrives from the periphery through the lemniscal thalamus to granular layers at 25ms and then propagates directly to the supragranular and infragranular layers (Figure 3A, left). The primary net effect of this input is synaptic activation of L2/3 interneurons and L2/3 pyramidal neurons through synapses on their basal and oblique dendrites. At 70ms, excitatory feedback projections — likely originating in SII (29) — arrive at supragranular targets, activating L2/3 interneurons and L2/3 and L5 pyramidal neuron tufts. At 135ms, a second, stronger “feedforward” input is presumed to arrive as part of an induced thalamocortical loop of activity. The raw SI current dipole response and net spiking activity in each cell population during this sequence of external perturbations are shown in Figure 3B, where black arrows mark the times of the exogenous drive. A smoothed version compared to the MEG measured SI evoked response are shown in Figure 3C/D.

**Figure 3:**
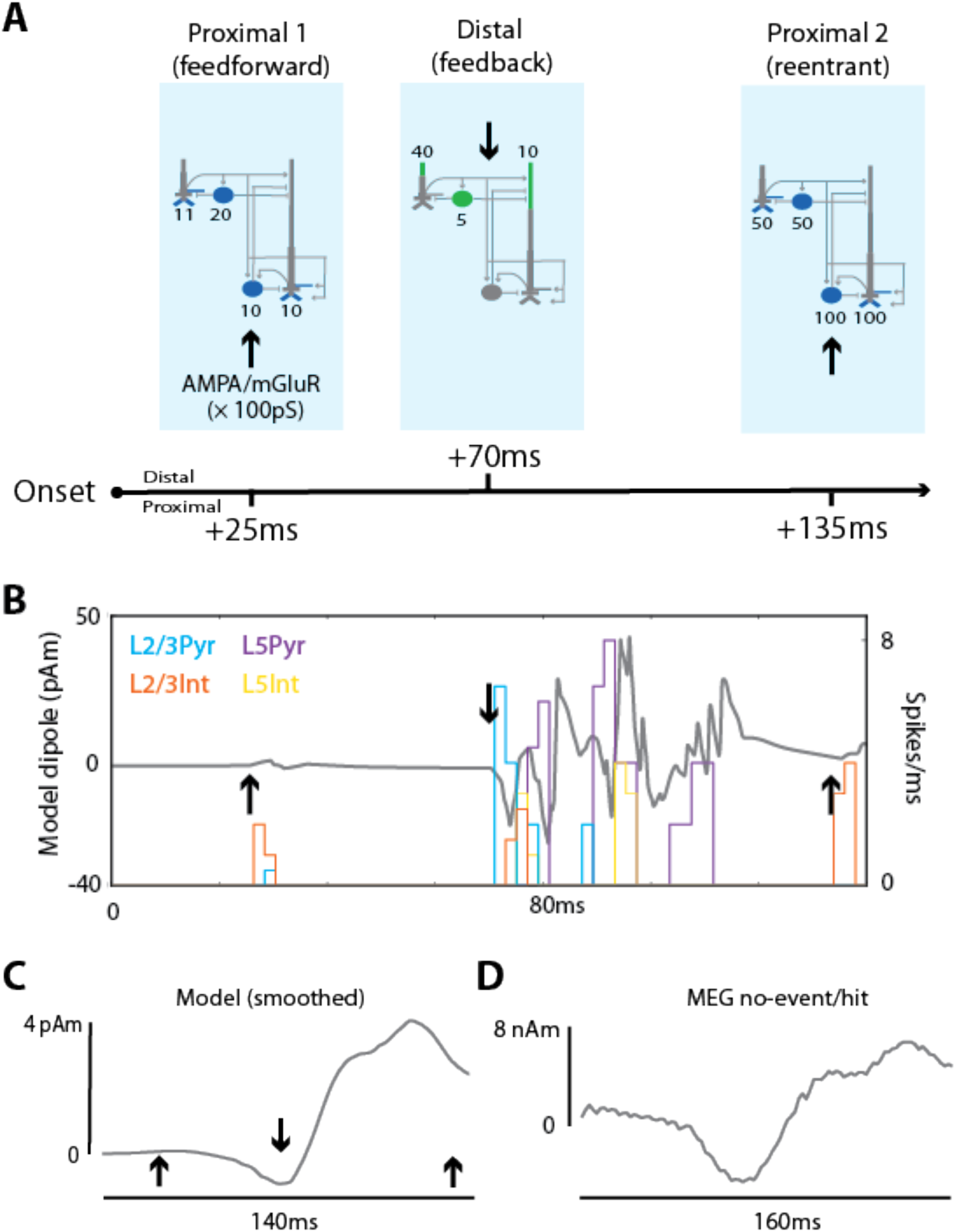
The SI model qualitatively reproduces the MEG-measured SI no-event/hit evoked response. **A)** Schematic of the feedforward/feedback exogenous input sequence reproducing the somatosensory evoked response. Sites of proximal glutamatergic “bottom-up” input are shown in blue and sites of distal “top-down” inputs in green. Pyramidal cells have three proximal sites (two on the basal dendrites, one on the oblique dendrite), each with the weight indicated here. **B)** Spike histograms and raw sensory evoked response from the model at rest (i.e. without latent currents from prestimulus activity): arrows same as in (A). **C)** The same model evoked response after convolving with a 45ms Hamming window. **D)** MEG evoked response averaged over all hit trials without a prestimulus beta event.

### Neurogliaform recruitment during beta generation is consistent with beta event waveshapes

We have previously shown that high power SI beta events have a distinct waveshape (10, see also highlighted waveform in Figure 1A). This shape was reproduced in our SI model by simultaneous subthreshold excitatory synaptic drive to proximal and distal pyramidal dendrites. As depicted in Figure 4B, a broad (~100ms) burst of subthreshold proximal drive pushes current flow up the pyramidal neuron dendrites, and a simultaneous, stronger distal burst pushes current flow down the dendrites for ~50ms (red arrow). The most prominent feature of the waveform is a downward deflection caused by the strong burst of distal drive, and lasting approximately one beta period (~50ms).

**Figure 4:**
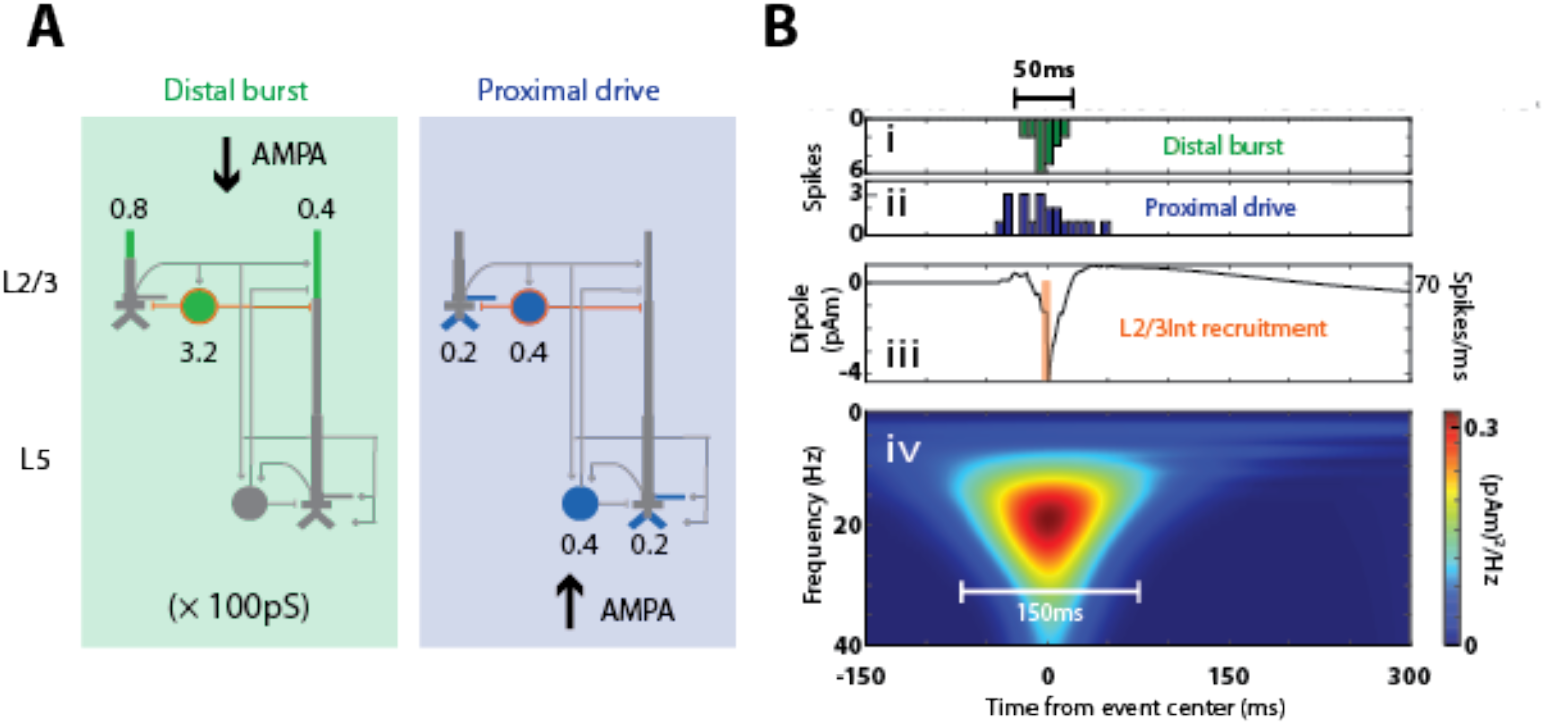
Simulating a beta event. Inputs to the model, modified from (Jones et al, 2009) during a simulated beta event (see Methods). **B) i.** Histogram of spikes incident on the proximal (blue) and distal (**ii**, green) dendrites of the cortical pyramidal and inhibitory neurons, as shown in (A), which generate the simulated beta event. Note that the beta event and evoked response input sites (Figure 3) are indistinguishable in the model. **iii** Corresponding current dipoles with concurrent spike histogram. Only the model L2/3 inhibitory neurons fire action potentials during a beta event. Activation of these interneurons causes GABA_B1a_ inhibition on pyramidal neurons, as shown in (A; orange); see also Experimental Procudures. **iv.** Time-frequency representation of the simulated beta event.

In the current study, we made a critical assumption based on the known circuit geometry of primary sensory cortex: we assume that the strong distal burst that generates the downward deflection of the beta event (Figure 4A/B) also recruits suprathreshold activation of a class of neurogliaform inhibitory cells in L2/3 (see Experimental Procedures, Beta Event Model). These cells activate slow metabotropic potassium channels (GIRK GABA_B_ channels), affecting L2/3 pyramidal neuron somata and L5 mid-apical dendrites (see Figure 4B).

To support this assumption, it was necessary to ascertain that the beta event waveshape (10) is preserved after NGF recruitment. A schematic showing synaptic weights onto the SI network in the NGF beta-event model is shown in Figure 4A with a model beta event shown in Figure 4B (see also Experimental Procedures). Stronger distal activation of L2/3 inhibitory neurons (Table 1) caused them to spike, while the remainder of the cells remained subthreshold.

The beta event waveform shape was largely maintained, even though L2/3 neurogliaform recruitment had distinct effects on the net current dipole. GABA_B_ slowly hyperpolarizes L2/3 somatic and L5 apical compartments with opposing effects, leading to downward dipoles from L2/3 and upward dipoles from L5 (data not shown). The summary effect of NGF recruitment on waveform shape was a long dipole tail after the beta event (see 250-500ms in Figure 4B) with power in the theta/delta bands but primarily concentrated near 20Hz as in our data.

### Model reproduces the influence of beta events on tactile evoked response via multiple latency-dependent effects

Having confirmed that the beta event waveform shape is preserved under our new modeling assumption, we next investigated beta event effects on evoked responses by simulating an evoked response (as in Figure 3) during and after a beta event (Figure 4). Our prior work shows that aggregate events are nearly uniformly distributed across the 1000ms prestimulus period (5). We inferred that some events were also likely to occur in the peristimulus interval before stimulus arrival and beta desynchronization at ~25ms, and therefore modeled beta event centers as occurring between −975ms to 25ms at 100ms increments.

There were multiple latency-dependent effects that — when averaged and compared to no-event evoked responses — reproduced beta’s mean effect on the MEG evoked response (Figure 5). This result emerged from the latency dependent impact of beta on the evoked response, without varying the evoked response inputs at each latency, and allows us to interpret beta dependent evoked response differences at the microcircuit level. Model parameters were tuned so that simulated evoked responses averaged over all latencies, when compared to no-event evoked responses, reproduced the primary differences observed in experimentally acquired SI MEG data (compare Figure 5D/E with F/G; statistical analysis was the same for model and data, see Experimental Procedures). After this qualitative fit was achieved, all model parameters were fixed — we did not alter the model to fit the predictions made in subsequent sections.

**Figure 5.**
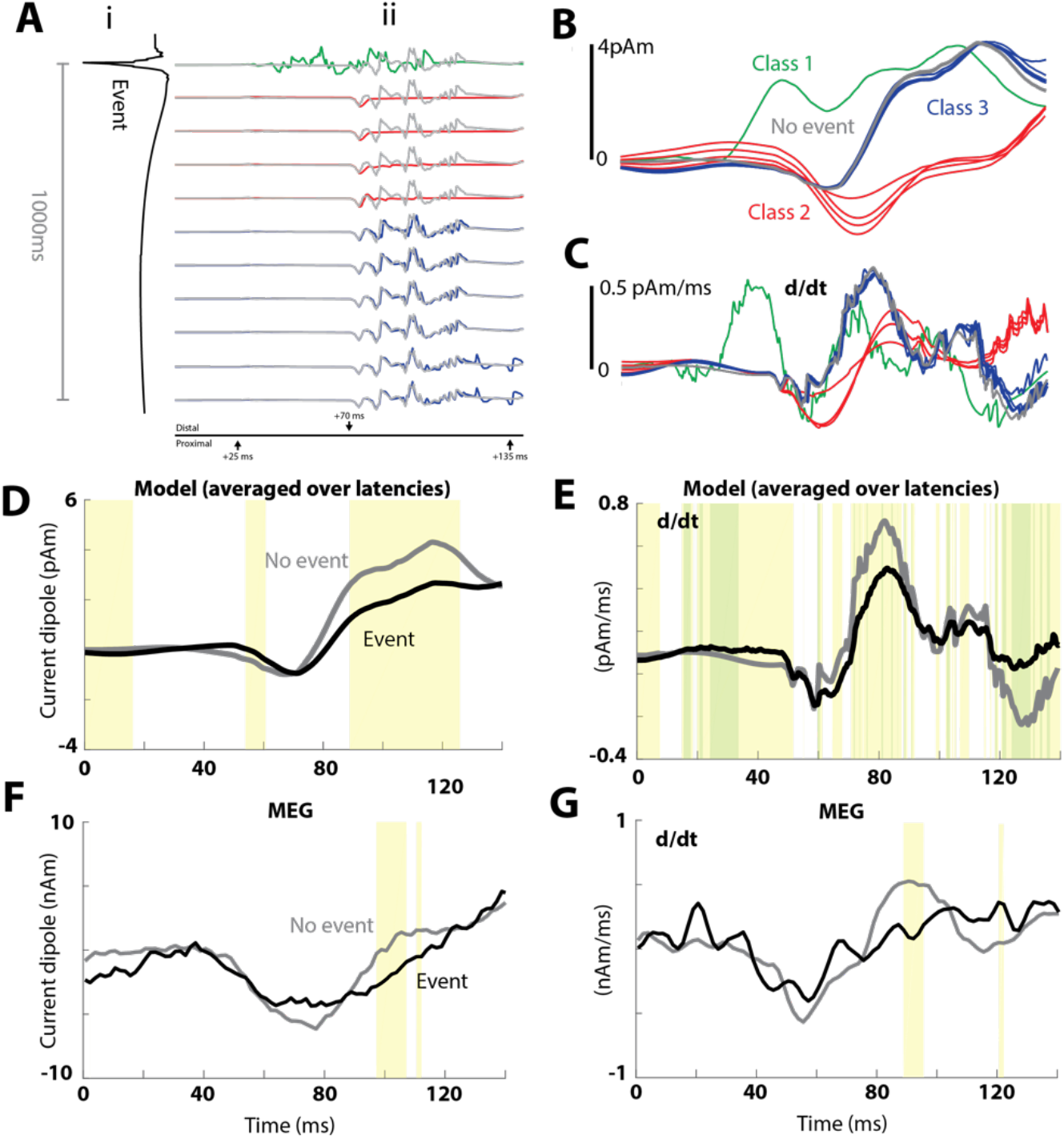
The SI model reproduces the impact of beta events on MEG evoked responses by averaging three latency-dependent response classes. **A)** Effects of beta events on model evoked responses depend on latency from the event. I) Beta event depicted from event onset to 1000ms after event onset (otherwise identical to Figure 3C). **ii)** Raw model evoked responses as a function of latency from event onset. (i) and (ii) are aligned such that (i) is the dipole during and following the event and (ii) is the evoked response when the first sensory input arrives in SI at the timepoint shown in (i). This +25ms shift accounts for conduction delay from the periphery. Time axis in panel ii is the same as in Figure 3A/B. Evoked responses from rest (see also Figure 3) are shown in grey for comparison. Color-coding represents a by-eye classification of response types. These three latency-dependent patterns are subsequently referred to as Class 1, Class 2 and Class 3 evoked responses. **B)** Smoothed model evoked responses, corresponding to the 11 raw latency-dependent responses in (Aii). **C)** Time derivatives of these smoothed responses, again color coded by class as in (Aii,B). **D)** Model evoked responses averaged over the 11 latencies in (Aii, B) compared to the evoked response from rest**. E)** Model time derivatives averaged over these latencies, as in (C). **F)** Human MEG SI evoked responses averaged over all trials with or without a beta event in the prestimulus period (reproduced from Figure 2A), and (G) their corresponding mean time-derivatives. Significance is reported both pointwise in yellow (p < 0.05; permutation test) and after FWER-correction in green (α = 0.05; modified max-T test; see Methods). Identical tests are applied to model and data.

The model generated three different evoked response patterns depending on latency from the beta event to stimulus onset, as color coded in Figure 5Aii: Class 1 (green), Class 2 (red) or Class 3 (blue). Class 1 responses occurred when the beta event trough and stimulus arrival (feedforward input) in SI coincide. During Class 1 responses, when the distally-incident burst and tactile inputs coincided, the model generated more prominent early dipole activity (~40 ms) than at other latencies. Class 2 responses occurred when the stimulus arrived between ~50ms-400ms after an event trough. Here, the initial dipole response to the feedforward input is nearly flat, and the dipoles were characterized by a unitary downward deflection at the time of the 70ms feedback input. Class 3 responses occurred when the stimulus arrived >400ms post-beta-event. These evoked responses were comparable to the no-event responses shown in gray, indicating that the effect of the beta event had worn off after 400ms. This is in agreement with our prior study showing that events tend to exert a strong bias toward nonperception when they occur 100–300ms preceding a stimulus (5). Smoothed responses and their time-derivatives are plotted in Figure 5B,C.

### Model evoked response classes are associated with distinct spike patterns that subserve correlates of detection

We can now address the important question: What are the precise circuit mechanisms underlying these beta event latency-dependent differences, and how might they relate to detection? To address this question, we examined model spiking activity for each cell type in our model during each class of response, and then inferred how these responses might generate correlates of detection in our MEG data.

Figure 6A shows model spike histograms along with raw (lighter) and smoothed (darker) net current dipoles. Layer specific responses are shown in panels B (L2/3) and C (L5). For all panels, spike rates and dipoles are averaged over all latencies (here, we modeled 101 latencies from −975–25ms at 10ms spacing) for a given class. In Class 1 responses (Figure 6Ai, 6Bi, 6Ci), feedforward sensory input arriving during a beta event trough recruits populations of L2/3 pyramidal cells to spike due to coincident depolarization from beta-event excitation of L2/3 pyramidal dendritic tufts (see Figure 4A) and excitation of basal and oblique dendrites from the stimulus-driven feedforward inputs (see Figure 3A). This “beta-primed” pattern is marked by early spiking activity in all L2/3 cells (Figure 6Bi) that spreads to spiking activity in all L5 cells, where the spiking persists for a longer period. These spike responses also created high frequency stimulus-locked dipole signals in L2/3, discussed further below.

**Figure 6:**
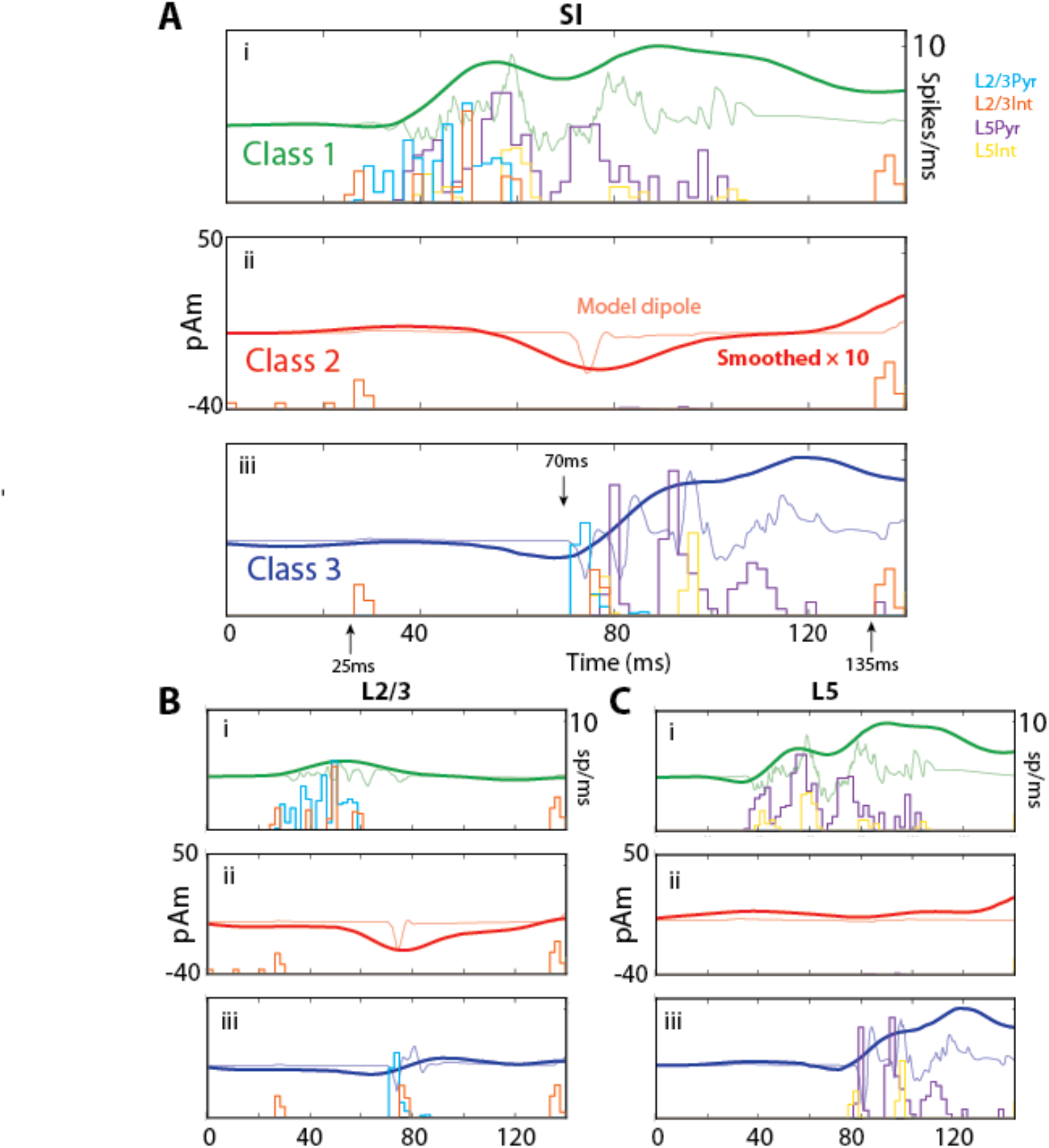
Evoked response classes correspond to spike-pattern classes with layer-specific generators. **A) i.** Class 1 evoked response averaged over each layer for 101 latencies (at 10ms rather than the 100ms resolution in Figure 5) and corresponding spike histograms for each cell type in the model. **ii.**and **iii)**. Same as (i), but for Class 2 and Class 3 responses. The Class 1 response shows early and persistent spiking, while the Class 2 response shows suppressed spiking. **B and C)** Contributions to the net current dipole response from L2/3 and L5 separately. **(i)** During the Class 1 response, L2/3 contributes to the early dipole and L5 to the later dipole**. (ii)** During Class 2 responses, L2/3 shows a prominent dipole deflection despite suppression of action potentials in both layers. **(iii)** The Class 3 response is similar to the no beta response in Figure 3.

Class 2 trials (Figure 6Aii, Bii, and Cii) were characterized by a near-complete absence of spikes in pyramidal cells of both layers (although isolated doublets from single cells in L5 appeared between 80 and 100ms late in the suppression period). The only spiking response from the initial feedforward input was in L2/3 interneurons — a population that had already been activated by a beta event. Note that the spikes appearing near 0, 10, and 20ms before the ~25ms feedforward input in the histograms in Figure 6Bii are from beta events themselves. Prestimulus recruitment of these interneurons is responsible for preventing spiking activity in both L2/3 and L5 mainly via the GABA_B_ hyperpolarization induced by the prestimulus beta event.

Importantly, the flat response in L5 is primarily responsible for the decreased amplitude and time-derivative observed for beta event trials in the model average (Figure 5C,D). The remaining prominent downward dipole in L2/3 was accounted for by large downward currents induced by the feedback input in the absence of spiking. Both of these effects are treated in more detail in a later section.

Class 3 trials (Figure 6Aiii, Biii, Ciii) occurred at latencies where the inhibitory effect of a prestimulus beta event had effectively worn off (i.e. >400ms after the beta event), and waveforms were similar to no-beta trials (Figure 3) with two exceptions. First, even small amounts of somatic GABA_B_ inhibition remaining more than 450ms after the beta event prevented L2/3 pyramidal spikes from the initial feedforward input. Second, for very long latencies (>800ms post-event; see Figure 5A), some pyramidal activity persisted until 135ms.

### Relating beta dependent model evoked response dynamics to MEG correlates of detection

We have shown that our model-based mechanism of beta event generation was able to accurately reproduce the observed influence of beta events on the tactile evoked response in MEG when averaging over multiple latencies (Figure 5D-G). Recall that the differences in evoked response characteristics between event and no-event trials, namely in the magnitude of the evoked response near 100ms and in the response derivative near 90ms, were analogous to differences between detected and non-detected trials (Figure 2). As such, we can now use our neural model to infer detailed circuit mechanisms underlying these correlates of detection in our data.

Of particular note, only Class 1 and Class 2 evoked responses differ significantly from the no-event evoked response. They have vastly different waveforms and spiking patterns that distinctly contribute to the averaged evoked response (Figure 5B). A primary distinction is that Class 1 trials exhibited a high proportion of pyramidal neuron firing, while Class 2 trials exhibited a near-complete absence of pyramidal neuron firing. This distinction led us to the unanticipated hypothesis that these responses might differentially correlate with detection; a finding that would have otherwise be masked by averaging.

Since L5 pyramidal neuron spiking communicates sensory signals from SI to higher order cognitive areas, we assume that sufficient pyramidal neuron firing in SI corresponds to detection, leading to the following two specific predictions.

Model prediction 1: Class 2 evoked responses with low pyramidal neuron spiking in the post-event period should correspond to miss trials in the MEG data.

Model prediction 2: Class 1 evoked responses with high early pyramidal neuron spiking should correspond to hit trials, and should be biased to occur during the falling and trough phases of a beta event in the MEG data (phases 0 to ½ in Figure 5A and 9A).

We test these two predictions with more specific comparisons between simulated and experimentally acquired MEG evoked responses below.

### Testing Model Prediction 1: Low spiking Class 2 evoked responses correspond to miss trials in the post-event period

We compared simulated Class 2 evoked responses to averaged evoked responses on miss trials in MEG data (Figure 7A). Remarkably, the Class 2 model evoked response was in immediate visual agreement with the averaged evoked response on miss trials with a prestimulus beta event — but also when there wasn’t a prestimulus event — suggesting similar inhibitory mechanisms can occur in no-event cases (Figure 7A; 502 event trials, 1498 no-event trials). A linear regression of the model Class 2 response onto the mean response of miss trials revealed nearly identical regression coefficients for each condition and for the average (r^2^=0.74, 0.83, 0.86, respectively; corresponding scaling factors needed to match the data amplitude are also shown in Figure 7A). The model scaling factor is approximately 2000 in all cases; this suggests that approximately 200,000 pyramidal neurons in L2/3 (100 cells in the model multiplied by this factor) are needed to generate the Class 2 response (as L5 did not generate significant dipole activity; see Figure 6Bii).

**Figure 7:**
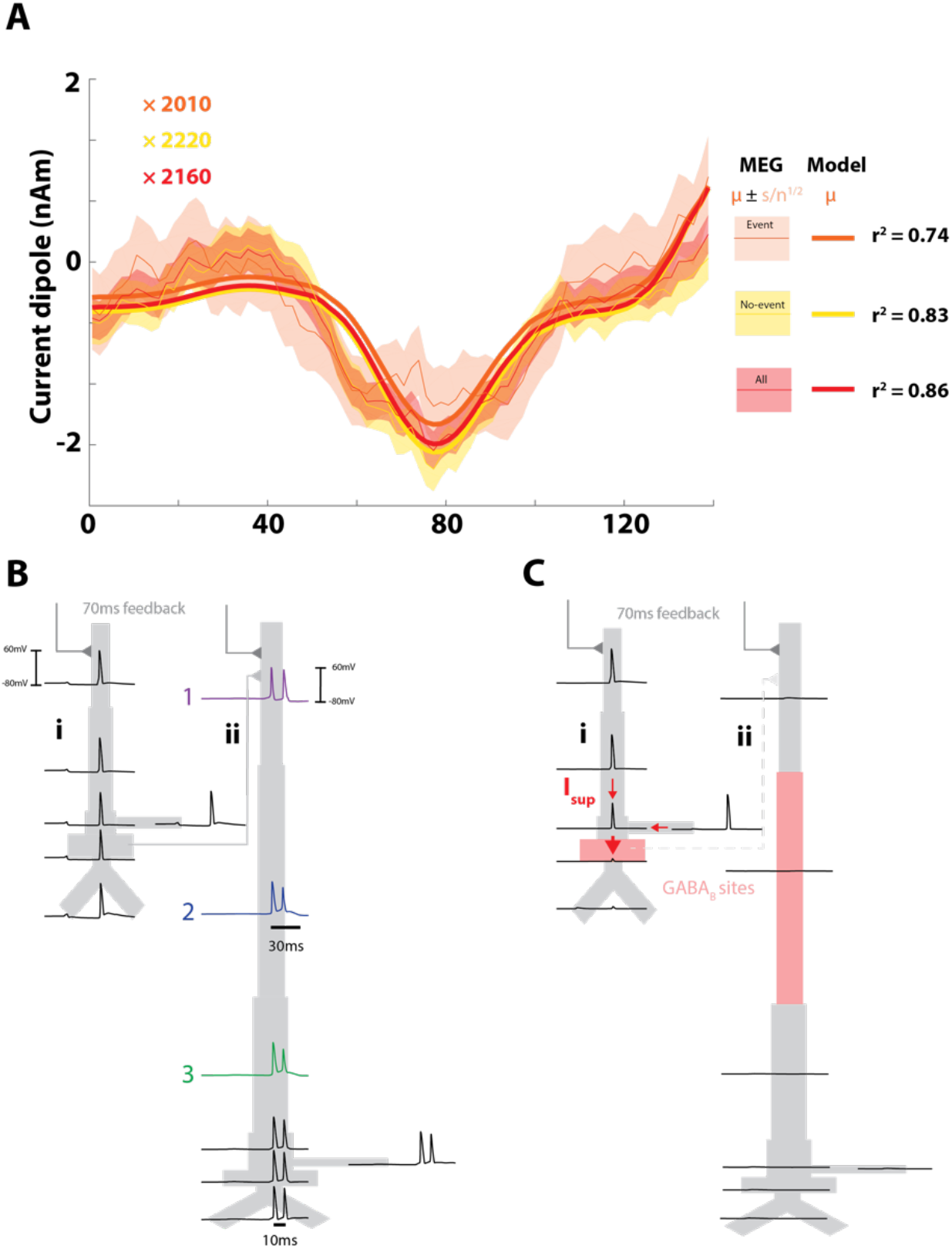
Model Class 2 responses are miss trials caused by active attenuation of dendritic spikes by strong perisomatic inhibition. **A.** The model Class 2 response directly correlates with the average MEG response for miss trials, whether or not a beta event was detected in the prestimulus period. The legend (right) indicates the model (thick lines) and MEG data distributions (thin lines; shading indicates standard error of the mean) for trials with and without events, and for all trials in aggregate. The model was fit by regressing directly onto the empirical means, and the corresponding r^2^ values and fitting parameters confirm a consistent agreement between model Class 2 trials and miss trials in MEG. Inset are scaling factors: multiplying these by 100 (the number of L2/3 pyramidal cells in the model) indicates that 200,000 L2/3 pyramids are sufficient to generate these dipoles. **B**. Membrane potentials are shown for each compartment of exemplar model pyramidal neurons within the same microcolumn, responding to “top-down” feedback arriving 70ms after stimulus onset. Without a prestimulus beta event (or other somatic inhibition; see Figure 5), both the L2/3 **(i)** and L 5 **(ii)** pyramidal neurons generate dendritic spikes that propagate to the soma after the top-down input. Indices 1-3 denote the temporal ordering for the first spike in the L5 burst based on inspection at higher temporal resolution (note that the second spike is initiated in the mid-apical dendrite rather than the tuft). **C)** After a prestimulus beta event, event-evoked GABA_B1a_ inhibition (i) prevents dendritic spikes from reaching the axon initial segment (considered as part of the somatic compartment) in L2/3 pyramids, and (ii) shunts the feedback response to L5, which prevents activity in L5 pyramidal neurons. Sites of inhibition are shaded pink; interneurons are not shown. Red arrows in **(Ci)** show that the difference in voltage between the soma and dendritic trunk in L2/3 pyramids creates an effectively maximal dipole current – extremized in both voltage gradient and conductive cross-sectional area—that generates the Class 2 evoked response shown in Figures 5B and 6.

Given that Class 2 responses mediate beta’s suppressive influence, we sought to more deeply understand the layer specific circuit mechanisms underlying this class of responses. Detailed cellular and circuit effects can be inferred by investigating the membrane potential in each compartment of example L2/3 and L5 pyramidal neurons at ~70ms post-stimulus during a simulated evoked response, with and without a prestimulus beta event (Figure 7B and 7C, respectively). We found that the observed lack of spiking in Layer 5 was due to a combination of processes; a lack of excitatory drive from L2/3 to L5 pyramidal neurons (compare L2/3 somatic responses in Figure 7Bi/Ci), and GABA_B_-mediated shunting of the response to the 70ms feedback input via inhibition at the mid-apical dendrites (compare response in L5 apical dendrites in Figure 7Bii/Cii; note, inhibitory spiking that provides GABA_B_ input (Figure 7Cii pink shaded regions) is shown in Figure 3). In comparison, L5 pyramidal neurons exhibit a calcium-mediated dendritic burst in response to the 70ms feedback drive in the absence of a prestimulus beta event. This burst starts in the distal dendrites and propagates to the soma: note the second spike in the burst is generated in the middle-apical dendrites (indicated with numbers 1-3 in Figure 7Bii).

To understand why the model Class 2 response exhibited a prominent ~70ms peak in the L 2/3 dipole waveform despite a lack of L2/3 spiking activity (Figure 6B, red), we again took a closer look at the difference between L2/3 cellular responses during no-event and post-event evoked responses (Figure 7Bi, 7Ci, respectively). Remarkably, the large dipole deflection for Class 2 responses was caused by downward dendritic spikes in L2/3 pyramidal cells that were blocked at the soma by GABA_B_ activity, which was not present in the no event case (compare somatic responses in Figure 7Bi/Ci). The voltage difference between the dendritic and somatic compartments creates a large intracellular dipole current in the Class 2 trials (see red arrows Fig 7Ci). This was indeed the largest local current allowed by the model neuron’s biophysics (see SI Appendix, Supplementary Calculation 1 for an estimate on thermal energy loss due to spike-blocking).

### Testing Model Prediction 2: High spiking Class 1 evoked responses correspond to hit trials and are biased to occur during falling and trough phases of a beta event

We expected Class 1 evoked responses to be rare in our MEG data, as they were not visible in hit trial averages (compare Figure 2B and Figure 6Ai) and because pre-stimulus beta events themselves were observed in only 25% of trials. However, additional phase coherence analyses (described below) allowed us to find Class 1 evoked responses (see Figure 8A), and to identify them as predominantly hit trials.

**Figure 8:**
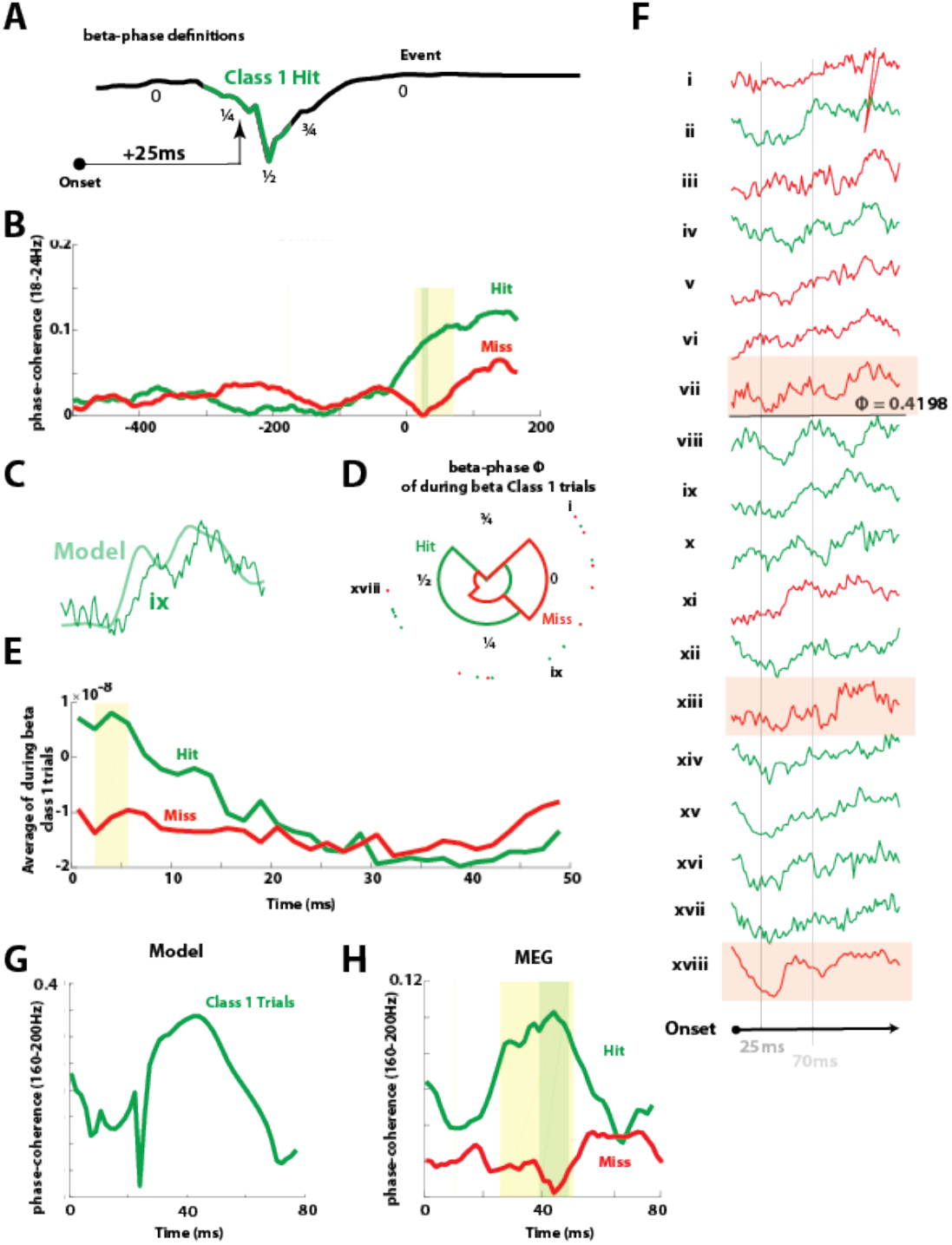
Class 1 trials are hit trials corresponding to event/stimulus matching. **A)** Beta event phases depicted on the model event. Green region depicts the approximate phase interval where Class 1 trials are predicted by the model (phase interval 0.25 – 0.75; normalized by 2𝜋) given the 25ms conduction delay from the periphery. **B)** Across-trial beta phase coherence (18-24Hz) in the MEG evoked response is shown for hit (green) and miss (red) trials. Poststimulus beta coherence is higher in hit trials, differentiating them at precisely 25ms poststimulus (TODO: include phase histogram with all trials as inset). **C)** The best-correlate between the model Class 1 response and empirical MEG evoked response is a hit trial. **D)** Beta phase at 25ms poststimulus nearly completely separates the Class 1 top-correlates into hit and miss trials (18 trials; 4 bins). Hit trials occurred within the phase interval predicted by the model with one exception. **E)** The eighteen top-correlates, averaged separately over hit and miss trials, show that hit trials exhibit an initial downward dipole slope on average. **F)** Direct inspection of all Class 1 top-correlates shows that 10 of 10 hit trials were either in dipole troughs or in downward phases at 25ms poststimulus, while 5 of 8 miss trials are in rising phases or plateaus. Extending the model-driven hypothesis to low-frequency activity outside the classical beta band accounts for this data nearly completely: Trials misclassified under this hypothesis are shaded pink. **G**) Across-trial high frequency coherence (160-200Hz) in model Class 1 responses predicts a peak in coherence near 40ms post-stimulus in hit trials. **H)** Across-trial 160–200Hz coherence distinguishes hit from miss trials near 40ms poststimulus (1000 hit trials, 1000 miss trials). In all panels, yellow shading indicates p < 0.05 (permutation test), and green shading indicates α = 0.05 (FWER-corrected; modified max-T test; see Methods). Significance in panel B is preserved at α = 0.02, and significance in panel H is preserved at α = 0.01.

First, the model predicted that during these rare Class 1 hit trials, the first feedforward sensory input to SI (at ~25ms post-stimulus;) coincides with the falling or trough phases of a beta event (Figure 8A; phase interval ~1/4-1/2). In order for this to be true, beta events would have to reliably occur during the post-stimulus period and be aligned such that their troughs occurred near ~25ms post-stimulus on at least a subset of hit trials. We tested this possibility by calculating beta phase coherence across trials in our MEG data (see Experimental Procedures, Figure 8B). We found that beta coherence in the post-stimulus period was significantly higher on hit compared to miss trials, with the highest significance precisely at ~25ms post-stimulus (22-32ms, alpha = 0.02; FWER-corrected permutation test). This suggests that at least some of the hit trials had a time locked beta event in the post-stimulus period.

Second, we sought to find Class 1 evoked responses in our data and to determine if those whose onset occurred during ¼ to ½-beta phase (corresponding to the falling or trough phases of a beta event, respectively; see Figure 8A) corresponded to hit trials. To find these responses, we performed a cross correlation analysis between the model Class 1 waveforms and our MEG trials, and then identified responses with the highest cross-correlation values between 0-25ms post-stimulus (see Experimental Procedures). We found 18 such Class 1 correlates at various possible phases of a poststimulus beta event in our data (Figure 8C-F). Figure 8C shows an example of the close agreement between the model and one of the identified Class 1 evoked responses in our MEG data. Approximately half of the identified Class 1 responses corresponded to hit trials (10 hit trials and 8 miss trials, green and red, respectively Figure 8F, corresponding phases are as in 8D). Averaging this subset of Class 1 trials separately over hit and miss trials revealed an important distinction between the responses, namely that the hit trials had a downward sloping dipole during the 0-25ms post stimulus period, while the miss trials did not (Figure 8E). This downward dipole deflection is consistent with the falling phases of a spontaneous beta event, as predicted by the model. Extending the phase histogram obtained through bandpass filtering, visual inspection indicates almost perfect separation between the hit and miss trials, such that all of the hit trials (green Figure 8F) indeed occurred during the falling and trough phases (10 of 10), as some of the underlying oscillatory events were outside the narrowband filter range. All except for 3 of 8 miss trials (red Figure 8F) occurred outside of the falling phases (pink highlight Figure 8F).

Third, we observed that during Class 1 evoked responses, the L2/3 pyramidal neurons in our model exhibited fast firing responses to the first feedforward input, creating a fast oscillation in the average dipole waveform beginning ~40ms (see raw green waveform in Figure 6Ai and 6Bi). The inter-peak interval in the dipole oscillation was approximately 6ms. As such, this observation led to the further model-based prediction that, if Class 1 responses exist and correspond to hit trials, then the Class 1 correlated hit trials in our data should have a reliable fast oscillating response in the dipole waveform with a frequency near 167Hz (i.e. 6ms period) that is not present in miss trials.

We quantified this prediction in our model by calculating high frequency coherence across trials (160-200Hz) for all Class 1 responses, and found a peak in cross-trial coherence around 40ms Figure 8G,H). We then tested if such high frequency coherence could be observed in the MEG data for hit but not miss trials, as predicted. We found higher coherence across trials on hit compared to miss trials; indeed, miss trial coherences were nearly zero throughout the early evoked response period (Figure 8H; alpha = 0.05; FWER-corrected permutation test). The timing of the significant differences between hit and miss trials in the MEG data were within 5ms precision of the times of peak coherence across trials in the model (40ms in the model, and approximately 40-45ms in MEG data).

**Figure 9:**
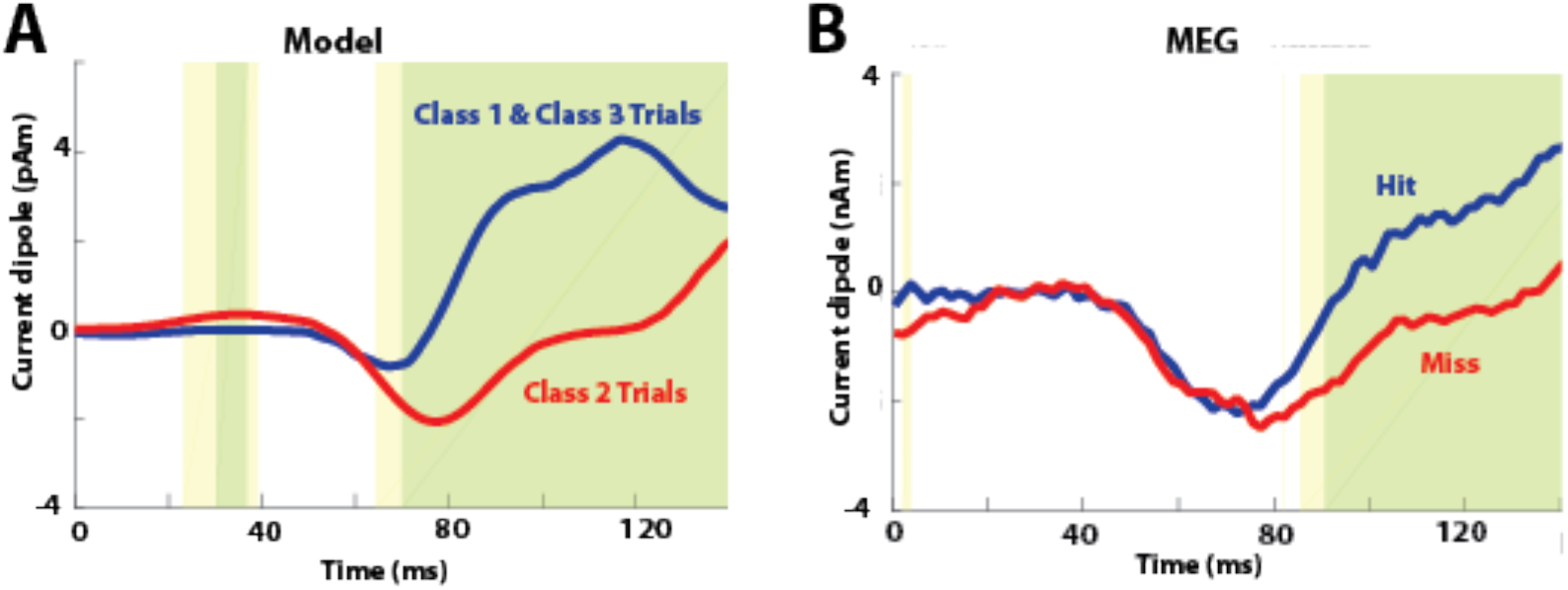
Classification by beta-event latency in the SI model reproduces the coarse dynamics of human tactile perception at threshold up to 140ms. **A)** The SI model of mean perceptual dynamics was conditioned on prestimulus beta events occurring with quasi-uniform (comb-distributed) timings. Model waveforms were first classified by latency (one Class 1 trial; four Class 2 trials; six class 3 trials, as in Figure 5), then assigned perceptual class membership (with Classes 1 and 3 as “hit” and Class 2 as “miss” trials, as in Figures 7–10). Means were then computed over each perceptual class. **B)** Mean perceptual dynamics in MEG (1000 hit, 1000 miss trials) source-localized to SI (yellow: p < 0.05; permutation test; green: FWER-corrected; α = 0.05; modified max-T test).

Finally, having established that Class 2 trials are predominantly miss trials, and Class 1 trials are predominantly hit trials, we assumed that Class 3 responses — analogous to no-event trials — would be mostly hit trials, as they exhibited strong spiking profiles. Comparing the latency-weighted average of Class 1 and Class 3 trials (all presumed-hit trials), to class 2 trials (all presumed-miss trials), we found clear agreement between model and MEG detection differences (Figure 10).

**Figure 10.**
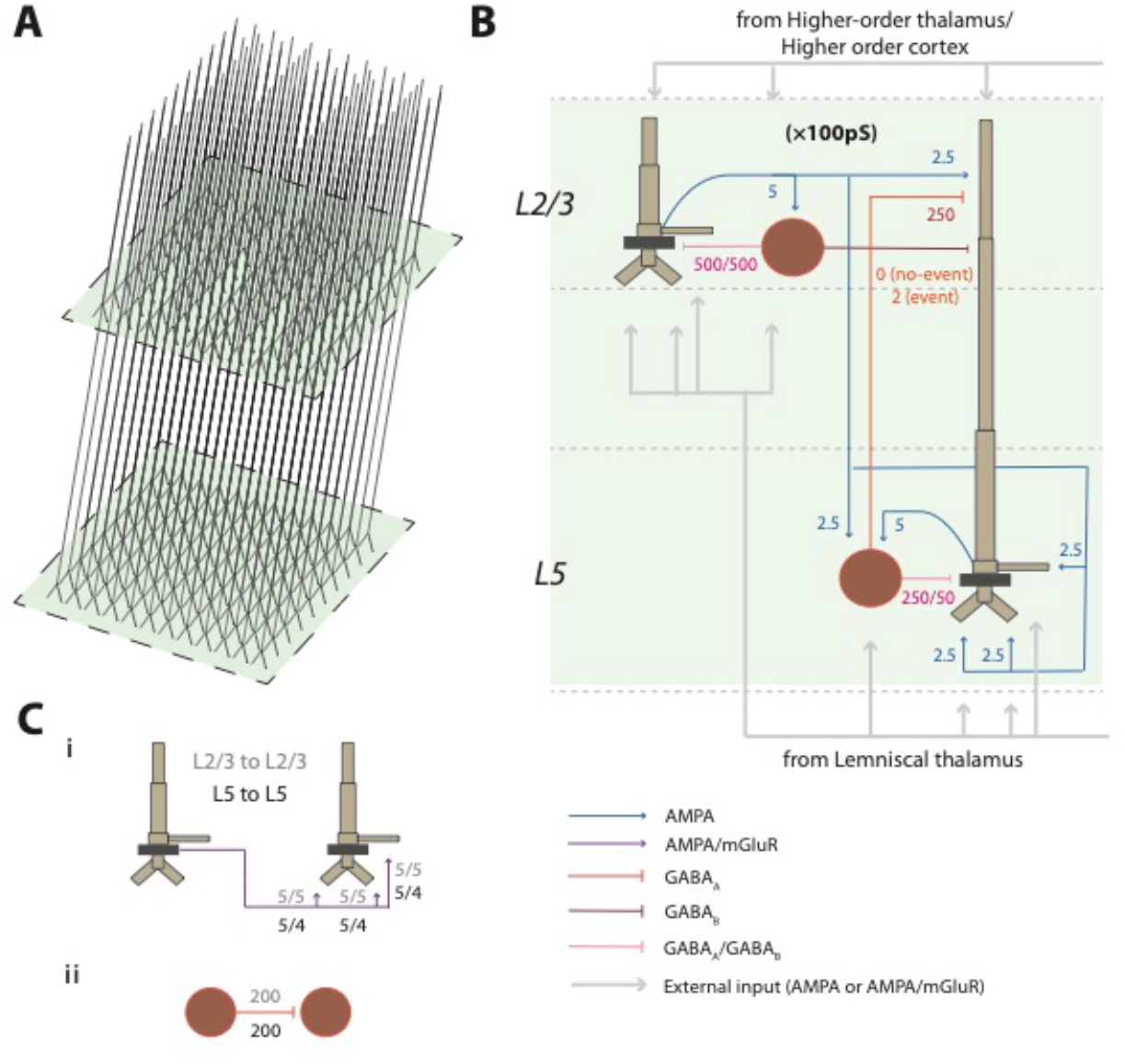
SI computational neural model. **A)** Model 3D pyramidal geometry shown approximately to scale. **B)** Reduced circuit schematic, with somata indicated in darker brown. Maximal conductances for each synaptic site are given in multiples of 10^-5 nS/cm^2. **C) i)** Schematic of lateral connections between pyramidal cells with synaptic weights labeled. **ii)** Schematic of lateral connections among interneurons.

## Discussion

Building from prior work showing that prestimulus beta events predict nondetection in a latency-dependent manner (5), we set out to understand the cellular and circuit level mechanisms by which this sensory suppression may occur in somatosensory cortex. Biophysically principled neural modeling combined with analysis of human MEG pointed to a parsimonious and quantitative explanation for this suppression: Beta events — thought to be electromagnetic signatures of higher-order thalamic bursts incident on cortex — activate supragranular pyramidal GABA_B1a_ receptors. This suppresses spiking throughout the cortical circuit, inhibiting sensory perception for ~400ms after the beta event.

In modeling latency, the model also showed that when a “top-down” beta event and tactile stimulus arrive in SI at the same time (a *coincident* or *on-stimulus* beta event), the coincidence generates an early cascade of L2/3 spiking, followed by L5 pyramidal bursting. This cascade created predicted MEG features that were verified as correlates of perceived trials by two separate means of testing (Figure 8). This result requires an interpretation of beta’s role beyond pure inhibition, as it indicates a mechanism by which beta can facilitate perception when timed appropriately.

### Consistency of the current findings with prior studies of SI evoked correlates of perception

It is well known that features of sensory-evoked responses can correlate with successful perception, and that spontaneous dynamic brain states, often described in terms of prestimulus “oscillations”, influence this correlation (20, 21, 30). However, an understanding of the precise neural mechanisms that could provide a causal link between prestimulus brain states and evoked measures of perception is lacking, particularly in humans where invasive recordings are rare. In the present study, our modeling framework enabled us to infer cellular and circuit-level mechanisms underlying the macroscale neural dynamic effects of beta events on the well-established tactile evoked response via direct comparison of model simulated data with source-localized MEG signals (same units of measure, Am). In so doing, we argue for a causal relationship between transient beta events and tactile detection, providing a detailed dissection of model processes ostensibly underlying the observed neural dynamics.

A critical model extension in the current study (10, 20, see also SI Appendix, Supplementary Discussion) was based on the novel hypothesis that higher-order thalamic mediated transient beta events recruit slow GABA_B1a_-mediated inhibition in the prestimulus time period. Updating our prior model to reflect this hypothesis did not significantly change the robustly observed stereotypical beta event waveform shape (10, 12), and was crucial to our conclusions regarding beta event-mediated tactile suppression. Our findings are also in agreement with several studies that have suggested slow inhibition in the supragranular layers — including through GABA_B_ mechanisms — is a key regulator of conscious perception (31, 32) and is specifically involved in inhibiting somatosensory perception (33). For example, Larkum and colleagues have shown that dendrite-mediated suppression of rodent somatosensory perception is regulated by GABA_B_ receptor activation in the mid-apical dendrite of L5 pyramidal cells, driven by a transcallosal inhibitory projection (28, 34). Our results extend theirs by modeling a local circuit mechanism for this inhibition in humans, noteworthy because long-range inhibitory projections are prevalent in rodents but have no known analog in primates (35).

The agreement between model output and MEG data support a causal influence of transient beta events on evoked correlates of perception, but it does not account for potential post-stimulus dynamics that are independent of beta events in the prestimulus period. In a previous study (21), we showed decreasing the arrival time and increasing the strength of the post-stimulus ~70ms distal “feedback” and subsequent ~135ms “re-entrant” thalamic inputs to SI (Figure 3) could account for detected evoked responses. In light of our current findings, we do not rule out the possibility of post-stimulus effects contributing to the evoked response on detected trials, where beta events are less likely to occur, and hence may also contribute to evoked correlates of perception.

### Anticipatory beta modulation could represent a predictive signaling mechanism

Our results show that SI beta events can both inhibit and enhance threshold-level tactile perception in a corresponding somatotopic region. Specifically, our findings suggest that if a beta event coincides precisely with tactile information, perception is enhanced, but after this time window closes, information processing is suppressed by slow inhibitory currents. Although on-stimulus beta events were rare in our data, this dichotomy raises an important conceptual debate*: Is beta actively engaged with task demands to inhibit perception? Or, might beta reflect a timing prediction mechanism with limited time windows that enhance perception, such that inhibitory prestimulus beta simply represents mis-timed (too early) beta events?* Our study was not designed to definitively address the cognitive strategy by which beta is temporally engaged. Below we discuss evidence in support of each possibility, see also SI Appendix, Supplemental Discussion where we discuss a learning mechanism that may be facilitated by beta suppression in order to optimize beta’s influence on perception.

Many studies have shown that cortical beta can be deployed with temporal specificity in coordination with task and cognitive demands (12, 19, 36–39). In the sensorimotor domain, we and others have observed that beta power decreases in localized somatotopic brain regions during the anticipatory period following a cue to attend to a corresponding body location, and increases in somatotopic regions corresponding to non-attended body locations (37, 40, 41). Such beta power modulation is also synchronized with frontal cortex (36), and peripheral muscles with or without task-specific motor demands (42). Lower prestimulus/anticipatory beta power leads to faster responses (37), and higher detection rates (5, 41), phenomena that have also been observed in visual-motion detection tasks where beta appears to track evidence accumulation (43). Moreover, it has been shown that beta power is actively decreased in a temporally specific manner, when a tactile stimulus is expected but not delivered (37). These and other studies (see also studies of motor cortex beta oscillations, where decreased beta activity corresponds with more certain responses and faster reaction times (12)) suggest sensorimotor beta is inhibitory to function and actively decreased for enhanced somatosensory detection, and/or actively increased to block irrelevant information.

Importantly, the data presented here was from a pure detection task, where the timing of the stimulus was randomized over 100ms intervals, and thus not easily predicted, and attention was not selectively manipulated or monitored. As such, in our task, prestimulus beta events may represent times when the subject’s “attention” strayed away from the finger, resulting in a corresponding increase in beta activity, supporting a sensory blocking strategy. Alternatively, the brain may be trying to engage beta at the “predicted” time of the stimulus in order to facilitate perception, such that prestimulus beta events represent premature timing predictions that instead suppress perception. It is worth noting that the aforementioned studies showing an inverse relation between beta power and perceptibility/motor action do not report beta phase. It is possible that smaller-amplitude events, with the appropriate phase alignment, are sufficient to facilitate processing when a prediction can be precisely made.

Our hypothesis regarding beta-mediated enhancement of perception was difficult to assess because beta power naturally decreases after sensory stimulation, a robust phenomenon known as beta event related desynchronization (ERD). However, several factors helped validate the existence of *on-stimulus* beta events during detected trials in our study, as our analyses suggest precise electrophysiological mechanisms and corresponding data features that can be used to identify such trials. Our MEG data show that perceived trials have falling low-frequency (in or near the beta band) phases at the time of stimulus arrival in SI. According to our proposed mechanism of the origin of beta events, these downward phases reflect ongoing excitatory bursts to apical dendrites of L2/3 pyramidal neurons, indicating a “top-down” signal inducing current flow toward the soma. This raises the potential of L2/3 pyramidal somata through passive cable mechanisms, such that simultaneous stimulus-evoked “bottom-up” excitation on the basal dendrite causes spike doublets in L2/3. The stimulus-locking of these spike doublets causes high-frequency coherence in the MEG signal, providing markers of coincident beta events and sensory stimuli on detected trials.

These on-stimulus beta events therefore represent a coincidence of “top-down” excitatory drive to apical dendrites and “bottom-up” drive to basal dendrites in L2/3 pyramidal neurons, aligning with a matching principle posed as a necessary component of conscious perception itself (44). Our model suggests an explicit locus and electrophysiological mechanism for this abstract principle, further specifying that a coincident match should be followed by a time-locked increase in spiking activity in L2/3, which later initiates L5 bursts. L5 bursts have themselves been argued to link a percept globally across the cortex (28).

Theoretical considerations further indicate that a temporal matching mechanism should appear in concert with a learning mechanism that processes timing errors — this may suggest a role beyond sensory suppression for Class 2 model responses. Recall that close agreement between model Class 2 and MEG miss trials led us to conclude that supragranular GABA_B_ inhibition accounts for beta’s latent perceptual suppression by abolishing SI interpyramidal communication. Despite this, remarkably large dipole currents remain, due to layer 2/3 dendritic spikes that propagate toward the soma but are extinguished upon arrival by inhibition (Figure 7). Several lines of evidence suggest that this process may provide a substrate for engaging “one-shot” postsynaptic learning (see SI Appendix, Supplementary Discussion and citations therein), in line with recent work indicating a causal relationship between beta events and plasticity (45).

## Conclusion

Further studies are needed to show if and how beta is actively modulated to suppress, amplify, and/or to learn the timing of sensory and/or internally-generated signals. Our present work extends and refines a variety of previous studies showing that beta events are dynamically engaged to meet task demands. These results are the first to posit explicit circuit mechanisms by which the presence of beta events can lead either to perception or nonperception depending on their timing. The upstream sources responsible for beta events — in particular higher-order thalamic bursts — can modulate cortical activity to gate the perceptual process at threshold. The work here points to new and precise cortical circuit mechanisms that can mediate this gating.

## Experimental Procedures

### MEG data collection and analysis

We analyze data from two previous studies, where MEG data localized to primary somatosensory cortex was collected during a tactile detection task. Detailed MEG and behavioral data collection methods can be found (20, 21), and relevant details and data analysis specific to the current study are described in (SI Appendix, MEG Data Collection and Analysis).

### Computational Neural Model Construction and Analysis

#### SI circuit model

We adapted the model structure from Jones et al. 2009 to reflect additional anatomical and physiological detail. Several changes to the 2009 model’s free parameters led to improved data fits and were informed by use of the Human Neocortical Neurosolver software (23). HNN freely distributes all of the code from the 2009 model, from which the current parameters were adapted. A description of differences between Jones et al. 2009 and the current model is in SI Appendix, SI Computational Neural Model Construction and Analysis and a direct comparison between parameters in each study are in Supplementary Table 1.

In brief, the model represents a layered cortical column with multicompartment pyramidal neurons and single compartment inhibitory neurons in supragranular layers 2/3 (L2/3) and infragranular layer 5 (L5), synaptically coupled with glutamatergic and GABAergic synapses (Figure 10). The primary current dipole signal is simulated by net intracellular current flow along the pyramidal axis in the direction of the voltage gradient between compartments following prior work (20, 21, 23). These currents are generated by active and passive membrane properties and synaptic dynamics (see Supplemental Table 1). Synaptic strengths in the current model are detailed in Figures 3, 4 and 10, and all other parameters used are detailed in Supplemental Table 1.

Details of how the evoked responses and beta events were simulated in the model are described in the Results section (see Figure 3 and Figure 4). Further details, including justification of the hypothesized beta event recruitment of long-time scale supragranular inhibition are in SI Appendix, SI Computational Neural Model Construction and Analysis, and a discussion of modeling assumptions, limitations and independence are in (SI Appendix, Supplemental Discussion).

### Comparing MEG and Model Data

#### Model fitting

Model parameters were tuned to reproduce evoked differences identified between event and no-event cases and their time-derivatives (Figures 3 and 5). Post hoc model tuning was not performed for quantitative data fits or predictions in subsequent analyses. In most simulations, the raw amplitude values from the model are reported in units of pAm, while the MEG data are reported in standard source-localized units of nAm. Assuming that the MEG signal is produced by large populations of synchronous cortical pyramidal neurons, a scaling factor can be applied to the model data to estimate the size of the network that contributes to the recorded response, as in our prior studies (20, 21). The scaling factor was calculated during direct comparison between model and MEG evoked response waveforms in Figure 7A.

Statistical analysis of waveform differences across conditions are described in (SI Appendix, Comparing MEG and Model Data).

## Acknowledgements

We thank Professor Matt Harrison for a number of helpful discussions and explanations, and for providing MATLAB code for the permutation and modified max-T tests. We thank Chris Black for assistance with Figure 10. Support for this project was provided by the NIMH (R01MH106174), NIBIB (R01EB022889), and the Department of Veterans Affairs, Veterans Health Administration, Office of Research and Development, Rehabilitation, Research and Development Service (Project N9228-C).

## Footnotes

* Here, we use “event” rather than “burst” so as not to confuse events with their mechanistic generators, which we believe to be bursts in upstream sources. Our interpretation of events is as “sequences of bursts”, viz. “bursts of bursts” where the length of the sequence can be as low as one.

