## Supplementary Materials for "A supragranular nexus for the effects of neocortical beta events on human tactile perception"

### Supplemental Data

#### Supplemental Table 1

The model consists of four cell types, of which only the two pyramidal cell types have multiple compartments and contribute to the dipole. Layer 2/3 pyramidal cells consist of a soma, an apical trunk directly above the soma, followed by apical 1 and, most distally, the apical tuft. The apical oblique dendrite branches horizontally from the apical trunk. There are also three basal dendritic compartments, with basal 1 directly below the soma and basal 2 and basal 3 branching from basal 1 at an angle, for a total of 8 compartments. The setup of the layer 5 pyramidal neurons is the same except that there is an additional apical compartment, apical 2, situated between apical 1 and the apical tuft. See Experimental Procedures and Figure 10 for a diagram of cell morphologies.

**Cell parameters**

| *Cell type* | *Parameter* | *Jones 2009* | *Law 2019* |
|---|---|---|---|
| L2/3 Pyr | Soma length (μm) | 22.1 | 22.1 |
| | Soma diameter (μm) | 23.4 | 23.4 |
| | Soma capacitive density (μF/cm$^2$) | 0.6195 | 0.6195 |
| | Soma resistivity (Ω-cm) | 200 | 200 |
| | Dendrite capacitive density (μF/cm$^2$) | 0.6195 | 0.6195 |
| | Dendrite resistivity (Ω-cm) | 200 | 200 |
| | Apical trunk length (μm) | 59.5 | 59.5 |
| | Apical trunk diameter (μm) | 4.25 | 4.25 |
| | Apical 1 length (μm) | 306 | 306 |
| | Apical 1 diameter (μm) | 4.08 | 4.08 |
| | Apical tuft length (μm) | 238 | 238 |
| | Apical tuft diameter (μm) | 3.4 | 3.4 |
| | Apical oblique length (μm) | 340 | 340 |
| | Apical oblique diameter (μm) | 3.91 | 3.91 |
| | Basal 1 length (μm) | 85 | 85 |
| | Basal 1 diameter (μm) | 4.25 | 4.25 |
| | Basal 2 length (μm) | 255 | 255 |
| | Basal 2 diameter (μm) | 2.72 | 2.72 |
| | Basal 3 length (μm) | 255 | 255 |
| | Basal 3 diameter (μm) | 2.72 | 2.72 |
| | AMPA reversal (mV) | 0 | 0 |
| | AMPA rise time (ms) | 0.5 | 0.5 |
| | AMPA decay time (ms) | 5 | 5 |
| | NMDA reversal (mV) | 0 | 0 |
| | NMDA rise time (ms) | 1 | 1 |
| | NMDA decay time (ms) | 20 | 20 |
| | $GABA_A$ reversal (mV) | -80 | -80 |
| | $GABA_A$ rise time (ms) | 0.5 | 0.5 |
| | $GABA_A$ decay time (ms) | 5 | 5 |
| | $GABA_B$ reversal (mV) | -80 | -80 |
| | $GABA_B$ rise time (ms) | 1 | 45 |
| | $GABA_B$ decay time (ms) | 20 | 200 |
| | Soma voltage-gated $K^+$ channel density (S/cm$^2$) | 0.01 | 0.01 |
| | Soma $Na^+$ channel density (S/cm$^2$) | 0.18 | 0.18 |
| | Soma leak channel reversal (mV) | -65 | -65 |
| | Soma leak channel density (S/cm$^2$) | 0.0000426 | 0.0000426 |
| | Soma M-type $K^+$ channel density (pS/μm$^2$) | 250 | 250 |
| | Dendrite voltage-gated $K^+$ channel density (S/cm$^2$) | 0.01 | 0.01 |
| | Dendrite $Na^+$ channel density (S/cm$^2$) | 0.15 | 0.15 |
| | Dendrite leak channel reversal (mV) | -65 | -65 |
| | Dendrite leak channel density (S/cm$^2$) | 0.0000426 | 0.0000426 |
| | Dendrite M-type $K^+$ channel density (pS/μm$^2$) | 250 | 250 |

| | | | |
|---|---|---|---|
| L5 Pyr | Soma length (μm) | 39 | 39 |
| | Soma diameter (μm) | 28.9 | 28.9 |
| | Soma capacitive density (μF/cm$^2$) | 0.85 | 0.85 |
| | Soma resistivity (Ω-cm) | 200 | 200 |
| | Dendrite capacitive density (μF/cm$^2$) | 0.85 | 0.85 |
| | Dendrite resistivity (Ω-cm) | 200 | 200 |
| | Apical trunk length (μm) | 102 | 102 |
| | Apical trunk diameter (μm) | 10.2 | 10.2 |
| | Apical 1 length (μm) | 680 | 680 |
| | Apical 1 diameter (μm) | 7.48 | 7.48 |
| | Apical 2 length (μm) | 680 | 680 |
| | Apical 2 diameter (μm) | 4.93 | 4.93 |
| | Apical tuft length (μm) | 425 | 425 |
| | Apical tuft diameter (μm) | 3.4 | 3.4 |
| | Apical oblique length (μm) | 255 | 255 |
| | Apical oblique diameter (μm) | 5.1 | 5.1 |
| | Basal 1 length (μm) | 85 | 85 |
| | Basal 1 diameter (μm) | 6.8 | 6.8 |
| | Basal 2 length (μm) | 255 | 255 |
| | Basal 2 diameter (μm) | 8.5 | 8.5 |
| | Basal 3 length (μm) | 255 | 255 |
| | Basal 3 diameter (μm) | 8.5 | 8.5 |
| | AMPA reversal (mV) | 0 | 0 |
| | AMPA rise time (ms) | 0.5 | 0.5 |
| | AMPA decay time (ms) | 5 | 5 |
| | NMDA reversal (mV) | 0 | 0 |
| | NMDA rise time (ms) | 1 | 1 |
| | NMDA decay time (ms) | 20 | 20 |
| | $GABA_A$ reversal (mV) | -80 | -80 |
| | $GABA_A$ rise time (ms) | 0.5 | 0.5 |
| | $GABA_A$ decay time (ms) | 5 | 5 |
| | $GABA_B$ reversal (mV) | -80 | -80 |
| | $GABA_B$ rise time (ms) | 1 | 45 |
| | $GABA_B$ decay time (ms) | 20 | 200 |
| | Soma voltage-gated $K^+$ channel density (S/cm$^2$) | 0.01 | 0.01 |
| | Soma $Na^+$ channel density (S/cm$^2$) | 0.16 | 0.16 |
| | Soma leak channel reversal (mV) | -65 | -65 |
| | Soma leak channel density (S/cm$^2$) | 0.0000426 | 0.0000426 |
| | Soma $Ca^{2+}$ channel density (pS/μm$^2$) | 60 | 0 |
| | Soma $Ca^{2+}$ decay time (ms) | 20 | 20 |
| | Soma $Ca^{2+}$-dependent $K^+$ channel density (pS/μm$^2$) | 0.0002 | 0.0002 |
| | Soma M-type $K^+$ channel density (pS/μm$^2$) | 200 | 200 |
| | Soma T-type $Ca^{2+}$ channel density (S/cm$^2$) | 0.0002 | 0.0002 |
| | Soma HCN channel density (S/cm$^2$) | 0.000001 | 0.000001 |
| | Dendrite voltage-gated $K^+$ channel density (S/cm$^2$) | 0.01 | 0.01 |
| | Dendrite $Na^+$ channel density (S/cm$^2$) | 0.14 | 0.14 |
| | Dendrite leak channel reversal (mV) | -71 | -71 |
| | Dendrite leak channel density (S/cm$^2$) | 0.0000426 | 0.0000426 |
| | Dendrite $Ca^{2+}$ channel density (pS/μm$^2$) | 60 | 60* |
| | Dendrite $Ca^{2+}$ decay time (ms) | 20 | 20 |
| | Dendrite $Ca^{2+}$-dependent $K^+$ channel density (pS/μm$^2$) | 0.0002 | 0.0002 |
| | Dendrite M-type $K^+$ channel density (pS/μm$^2$) | 200 | 200 |
| | Dendrite T-type $Ca^{2+}$ channel density (S/cm$^2$) | 0.0002 | 0.0002 |
| | Dendrite HCN channel density (S/cm$^2$) | 0.000001** | 0.00001** |

* Density is equal to this value in apical dendrites (apical trunk, apical oblique, apical 1, apical 2, and apical tuft) only, and equal to 0 in the basal dendrites (basal 1, basal 2, basal 3).
** HCN channel density increases as an exponential function of distance from the soma, starting at 0.000001 S/cm$^2$ in the soma and increasing with a space constant of 0.003.

Figure 10 provides a schematic of the connectivity between cell types in the model. Within each layer, synaptic strengths between pyramidal neurons are scaled according to a 2D symmetric Gaussian defined on the grid of cells, with a weight space constant of 3. Similarly, the synaptic delay between two PNs in the same layer is scaled according to an inverse Gaussian with a delay space constant of 3.

**Network connectivity parameters**

| *Source cell* | *Target cell* | *Synapse* | *Maximal conductance (µS)* | |
|---|---|---|---|---|
| | | | *Jones 2009* | *Law 2019* |
| L2/3 Pyr | L2/3 Pyr | AMPA | 0.0005 | 0.0005 |
| | | NMDA | 0.0005 | 0.0005 |
| | L2/3 Basket | AMPA | 0.0005 | 0.0005 |
| | L5 Pyr | AMPA | 0.00025 | 0.00025 |
| | L5 Basket | AMPA | 0.00025 | 0.00025 |
| L2/3 Basket | L2/3 Pyr | $GABA_A$ | 0.05 | 0.05 |
| | | $GABA_B$ | 0.05 | 0.05 |
| | L2/3 Basket | $GABA_A$ | 0.02 | 0.02 |
| | L5 Pyr | $GABA_A$ | 0.001 | 0 |
| | | $GABA_B$ | 0 | 0.0002 |
| L5 Pyr | L5 Pyr | AMPA | 0.0005 | 0.0005 |
| | | NMDA | 0.0005 | 0.0004 |
| | L5 Basket | AMPA | 0.0005 | 0.0005 |
| L5 Basket | L5 Pyr | $GABA_A$ | 0.025 | 0.02 |
| | | $GABA_B$ | 0.025 | 0.005 |
| | L5 Basket | $GABA_A$ | 0.02 | 0.02 |

**ERP input parameters**

| *Input type* | *Target cell* | *Synapse* | *Maximal conductance (µS)* | |
|---|---|---|---|---|
| | | | *Jones 2009* | *Law 2019* |
| Early proximal | L2/3 Pyr | AMPA | 0.001 | 0.0011 |
| | L2/3 Basket | AMPA | 0.002 | 0.002 |
| | L5 Pyr | AMPA | 0.0005 | 0.001 |
| | L5 Basket | AMPA | 0.001 | 0.001 |
| Distal | L2/3 Pyr | AMPA | 0.001 | 0.004 |
| | | NMDA | 0.001 | 0.004 |
| | L2/3 Basket | AMPA | 0.0005 | 0.0005 |
| | | NMDA | 0.0005 | 0.0005 |
| | L5 Pyr | AMPA | 0.001 | 0.0005 |
| | | NMDA | 0.001 | 0.0005 |
| Late proximal | L2/3 Pyr | AMPA | 0.0053 | 0.005 |
| | L2/3 Basket | AMPA | 0.0053 | 0.005 |
| | L5 Pyr | AMPA | 0.0027 | 0.01 |
| | L5 Basket | AMPA | 0.0027 | 0.01 |

**Beta event parameters**

*Burst properties*

| *Input type* | *Parameter* | *Sherman 2016* | *Law 2019* |
|---|---|---|---|
| Proximal | Standard deviation (ms) | 20 | 20 |
| | Number of bursts | 10 | 10 |
| | Spikes per burst | 2 | 2 |
| | L2/3 delay (ms) | 0.1 | 0.1 |
| | L5 delay (ms) | 1.0 | 1.0 |
| Distal | Standard deviation (ms) | 10 | 10 |
| | Number of bursts | 10 | 10 |
| | Spikes per burst | 2 | 2 |
| | L2/3 delay (ms) | 5 | 0.5 |
| | L5 delay (ms) | 5 | 0.5 |

*AMPA conductances (µS)*

| *Input type* | *Target cell* | *Compartment* | *Sherman 2016* | *Law 2019* |
|---|---|---|---|---|
| Proximal | L2/3 Pyr | Basal 2 | 0.00002 | 0.00002 |
| | | Basal 3 | 0.00002 | 0.00002 |
| | | Apical oblique | 0.00002 | 0.00002 |
| | L2/3 Basket | Soma | 0.00004 | 0.00004 |
| | L5 Pyr | Basal 2 | 0.00002 | 0.00002 |
| | | Basal 3 | 0.00002 | 0.00002 |
| | | Apical oblique | 0.00002 | 0.00002 |
| | L5 Basket | Soma | 0.00002 | 0.00002 |
| Distal | L2/3 Pyr | Apical tuft | 0.00004 | 0.00008 |
| | L2/3 Basket | Soma | 0.00008 | 0.00032 |
| | L5 Pyr | Apical tuft | 0.00004 | 0.00004 |

### Supplemental Calculation 1: Spike destruction accompanies biophysically maximal dipoles

The model "Class 2" evoked response dominates nonperceived trial averages despite a generating population of no more than 200,000 model L2/3 pyramids (see Figure 7 and Results). Although dipoles scale with their spatial separation, this large current dipole is generated by a near point-source. We show here that it is indeed the largest current dipole that can be generated by the model cell.

We first show that the current dipole dependence on length disappears in a linear resistive medium, as in the far-field we can view the net current dipole (*D*) as a sum of dipoles from all compartments *c*.

$$D = \sum_c I_z \Delta z$$

Intracellular axial current flow is governed by $I = g(V_1 - V_0)$ where 1 and 0 represent upper and lower cable junctions (compartment boundaries), g the intracellular conductance, and V the transmembrane voltage (conventionally assuming uniformity of the extracellular fluid).

For each compartment, conductance scales inversely with length $\ell$ and directly with cross-sectional area $A$, which is assumed to be maximized in the soma. The one-compartment current dipole state is $D = \sigma^{-1} \frac{A}{\ell} \Delta V \Delta z$. Observing that $\Delta z = \ell$, the current dipole for any compartment is then

$$D_c = \frac{A_c}{\sigma_c}(V_1 - V_0)$$

This shows that given our assumptions, current dipoles are *independent of compartment length*. Geometrically, they depend on cross-sectional area instead; this is physically reasonable as magnetic fields depend on current flux density through a surface. Note that magnetic fields measured in MEG (fT) were converted to current dipole units (nAm) during the source localization procedure (1, 2).

The state model is now equivalent to a series composition of simple resistors where we may set junction voltages arbitrarily to find a configuration that maximizes the net directional current flux. The maximum current dipole occurs when all junctions above the soma are at $V_{max}$ and all junctions below the soma are at $V_{min}$. Here, $|\Delta V|, A$ are both maximized despite generator length being essentially minimized.

Conditions in a cell that maximize current dipoles will be said to satisfy the *extreme local current property*. In models of adult mammalian cells without calcium currents, see e.g. (3), membrane voltages are bounded by the respective sodium and potassium reversal potentials, i.e. $E_{Na} = V_{max}$ and $E_K = V_{min}$. Figure 7C confirms the extreme local current property holds approximately for sodium dendritic spikes where strong somatic inhibition (viewed as a control signal, denoted U) acts as a spike destructor function:

$$f: (U, spike) \to (U, 0)$$

**Supplemental Calculation 2: Estimated thermal activity due to spike destruction**

If an extremal current is indeed localized to a somatic compartment, the following estimate shows the thermal side effects of spike destruction are nonnegligible.

Energy conservation governs heating in simple electrical resistors, and if linear resistive models of cytoplasm suffice, the average thermal energy dissipated over a compartment (*Joule heating*) for a distorted spike is estimated as:

$$\Delta E = \Delta \text{t}(\Delta V)^2/R$$

The reader may verify that one destroyed spike generates thermal energy in the sub-nJ range for our model parameters, and one may also wish to estimate the energetic upper bound where spike destruction occurs simultaneously in all pyramidal somata.

Here, though, we are interested in the rate of temperature change in a volume $\mathcal{V}$ (the soma) containing a substance (water) with density $\rho$ (= $1 \times 10^{-6}$ μg/μm$^3$) and specific heat capacity $C$ $(= 4.186\ \mu J/(\mu g \cdot K))$ as a spike of amplitude $\Delta V (= 100 \text{ mV})$ is destroyed while traversing that volume.

Recalling $\Delta T = \frac{\Delta E}{mC} = \frac{\Delta E}{\rho \ell A C}$ for a linear compartment (where $\mathcal{V} = A\ell$ for cross-sectional area A) we have:

$$\frac{\Delta T}{\Delta t} = \frac{(\Delta V)^2}{\rho C \ell A R}$$

The resistance scales linearly with length (in series) and inversely with cross-sectional area (in parallel), with model resistivity $\sigma = 200\ \Omega\text{cm}$ (4, 5). Substituting $R = \sigma\ell/A$ into the above equation (while again noting that $\Delta z = \ell$) yields:

$$\frac{\Delta T}{\Delta t} = \frac{(\Delta V)^2}{\sigma\rho C(\Delta z)^2}$$

$\cong 2.5\ K/s$ ,

assuming the spike is destroyed over the model L2/3 pyramidal somatic compartment, a drum-shaped cable segment with height 22.1$\mu m$. Under alternate conditions ($150mV$ attenuation, $150\Omega cm$ resistivity), we obtain instead 7.5K/s. Large differences in TRPV3 cationic currents were reported in this range of thermalization (6).

Two exemplar studies exhibiting a remarkable range of thermodynamic processes in living cells can be found in (7, 8). We observe that the electrothermal shocks associated with spike destruction are inverse-quadratically sensitive to the axial length of the prosomatic "shock zone" wherein a spike is destroyed. If pronounced in real neurons, the effect will likely depend on the distribution of perisomatic inhibition – our model uses a point synapse at the middle of the somatic compartment. Absolute temperature changes will depend on the rate of destroyed spikes and the diffusion tensor.

### Supplemental Text Describing Experimental Procedures

#### MEG Data Collection and Analysis

*Tactile-detection task*

A 2kHz auditory cue was presented for a duration of 2000ms. The timing of each tactile stimulus was sampled uniformly from a comb distribution spanning 500–1500ms after cue onset, consisting of 11 evenly spaced postcue timings. The stimulus consisted of a single sinusoidal tactile pulse lasting 10ms, delivered from a piezoelectric ceramic benderplate to the tip of the D3 digit of the right hand. After cue offset, subjects were instructed to press a button with the left hand (second or third digit) to report if the stimulus was perceived or not perceived. Tactile stimulus pulse amplitude was initially tuned to the individual's perceptual threshold using the parameter estimation by sequential testing method (9, 10) and then dynamically maintained around threshold (1).

*MEG dipole source localization and preprocessing*

MEG dipoles were source-localized to primary somatosensory cortex using a two-dipole model (SI and SII), fitting suprathreshold responses with a signal-space projection method (11, 12). Data were sampled at 600Hz and bandpassed online between 0.01Hz and 200Hz. Individual trials were epoched into prestimulus (-1000ms – 0ms, where 0ms indicates tactile pulse onset) and poststimulus (0 – 2500ms) time periods. For each individual, the final 100 hit (detected) and 100 miss (non-detected) trials were isolated for further analysis due to the stability of the dynamics at the end of the experiment (13).

*Baseline normalization and time-derivative estimation*

The choice of baseline correction method has been shown to differentially bias bandlimited power in postcue periods (13), and it was more recently shown directly that oscillations are coupled to baseline changes (14). If events influence later responses, baseline correction may also introduce spurious biases that will confound modeling attempts. In our study, the downward-bias of beta event waveforms (15) consistently lowers event-trial dipole means, and compensatory shifting intended to correct for these prestimulus differences introduced strong but spurious differences in poststimulus dipoles.

To ensure that we did not fit our model to biased differences, we did not baseline shift our data. We examined both raw data and time-derivatives — these yield baseline-free comparisons of dipole changes from active membrane processes. We used total-variation regularized differentiation, which corrects noise amplification in first-difference estimates of time-derivatives while preserving jump discontinuities observed in MEG data without linear filtering artifacts (16). We fixed the regularization parameter at $10^{-8}$ based on visual inspection of regularized fits to two trials (not shown).

*Defining beta events and assessing their perceptual effects*

Events in the prestimulus period were identified as regions in the time-frequency plane whose energy crossed six times the median energy at a given frequency, computed separately for each individual by convolution with a six-cycle Morlet wavelet (see 7 for more detailed methods). In this study we restricted our analysis to events with energy peaks between 20-22Hz, following the spectral concentration within the beta wideband (15-29Hz) observed in human somatosensory recordings (2, 18). Trials were then partitioned into those with one or more prestimulus events (*event* trials) and those without events (*no-event* trials).

We used the Wilcoxon rank-sum test to assess whether narrowband beta events affected perception (see Figure 1B).

*Post-stimulus phase coherence*

For all post-stimulus phase-coherence analyses, we first filtered the data in the frequency band of interest across the entire trial [-1000, 2500ms] and then applied a Hilbert transform to define phase, after which phase coherence was computed across trials.

We obtained results pertaining to poststimulus phase coherence (Figure 8B, G and H) with a Chebyshev bandpass filter at 10dB within-band ripple. We initially chose this filter due to its rolloff properties useful for narrowband analysis, as higher-order Butterworth filters failed due to numerical issues. Because the Chebyshev filter amplifies at the bandlimits, we also determined whether a more conventional filter choice could yield similar results. Events with energy peaks at 20-22Hz had a nearly flat event-density distribution between 18–24Hz (data not shown), justifying the use of a third-order Butterworth filter in this wider band for beta phase analysis, and where we obtained nearly identical results.

For the 160-200Hz Chebyshev filter, we found that a third-order Butterworth filter in the 178-197Hz band also reproduced the significant differences between hit and miss trials at similar or identical timepoints. We report one result from each filter for beta- (3rd order Butterworth; 18–24Hz) and high-frequency phase (Chebyshev, 10dB within-band ripple; 160-200Hz).

After bandpassing, we computed the Hilbert transform for each trial $x$:

$$H(x) = \mathcal{F}^{-1}(2u \cdot \mathcal{F}(x))$$

where $\mathcal{F}$ is the (fast) Fourier transform and $u$ is the Heaviside step function. Real-valued sequences are paired with complex sequences, resulting in analytic signals $x + iH(x)$ with instantaneous phase given by the complex angle $\phi = \arctan \frac{H(x)}{x}$. For narrowband beta events, the trough is associated with phase $\pi$ —Note that under this convention phase decreases over time.

We then computed the cross-trial phase coherence at each timepoint as the modulus of the vector mean:

$$C = \frac{|\Sigma e^{i\phi(t)}|}{N}$$

Note that zero coherence does not imply a uniform distribution across phases, but rather that the sample is phase-cancellative across trials.

*Correlating Class 2 model waveforms and MEG miss trials*
To compare "Class 2" (in which a beta event precedes the tactile stimulus by ~100-400 ms) model waveforms to MEG miss trials (Figure 7A), the mean empirical evoked response waveform ($\mu_{MEG}$) was linearly regressed by least-squares directly onto the smoothed model waveform ($x_{model}$), obtaining:

$$\mu_{MEG} \approx \beta_0 + \beta_1 \cdot x_{model}$$

The $r^2$ values assess agreement between model and data. We performed regressions separately for miss trials in event and no-event cases as well as in aggregate. In all cases, the constant offset parameter $\beta_0$ was small and not considered further. The linear scaling parameter $\beta_1$ was similar in all cases and used to estimate the scaling factor applied to the model to estimate the number of neurons that can account for the magnitude of the recorded signal.

We report error regions reflecting standard error of the mean $\frac{s}{n^{1/2}}$ for reference in Figure 7A, but we do not report the standard error in other figures as our evidence of multiple waveform classes indicates that this statistic, which should only be interpreted under normality of residuals, would be misleading for interpreting our data.

*Finding direct and indirect correlates of Class 1 evoked responses in MEG data*

We expected Class 1 trials (in which a beta event and tactile stimulus occur at the same time) to be rare (see Results) making their identification sensitive to ongoing activity that might vary on a trial wise or individual basis. We furthermore did not want to exclude candidate trials due to small time-lags unaccounted for by our model. However, because Class 1 trials were primarily characterized by a dipole upswing near 40ms poststimulus (see Figures 6, 8), false-positives could be generated by unrelated high-amplitude transients or indeed any waveform with an upward trend.

To find candidate Class 1 trials in such a way that would be unbiased by within-trial variability, we first calculated the normalized cross-correlation $\chi(x_{model}, x_{MEG})$ between the model waveform and each MEG trial -- over the entire trial (-1000–2500ms) -- and found the lag time of peak cross-correlation for each trial i:

$$\tau_i = \text{argmax}\, \chi_i$$

Histograms indicated a large peak in the distribution of peak Class 1 cross-correlates near 50ms lag (data not shown), an indicator of spurious correlation with the 80-90ms upward deflection exhibited in a significant fraction of all trials. These trials are more likely to be Class 2 or Class 3 trials: We excluded these, as well as trials with peak cross-correlation before onset, restricting our analysis to candidate Class 1 trials with lags $0ms \leq \tau \leq 25ms$. This interval would not confound Class 1 with Class 3 trials, which exhibit similar waveforms but with different time-delays. This method also generated a sample small enough (n=18) where all waveforms could be presented, but large enough where statistics could be meaningfully computed. Similar results were obtained in wider intervals, up to $-5ms \leq \tau \leq 30ms$.

The model having indicated that beta phase at 25ms poststimulus should covary with detection, we then examined how this feature, in combination with cross-correlation, dissociated Class 1 candidates into hit and miss trials. To do so, we simply examined a circular histogram of beta phases for the 18 candidate Class 1 trials.

### Supplemental Text Describing Computational Model

#### Computational Neural Model Construction and Analysis

*Comparison between current model and Jones et al. 2009*

Cell dendrite and somatic compartments were simulated with active ionic currents, as detailed in our prior study (2), see Supplementary Table 1. Structural differences between the 2009 model and current model included addition of Martinotti-like recurrent tuft connections from the L5 interneuron to the L5 pyramidal neurons' distal dendrites and removal of L2/3 $GABA_A$ interneuron synapses onto those same dendrites. We also adapted the model's L5 calcium channel distribution, restricting expression to the apical dendrite. The most important *a priori* change in the model was an increase of the $GABA_B$ channel time constants to reflect data reported in (19). We reduced this model to a double-exponential conductance with 45/200ms respective rise/fall times, matching the peak conductance latency ~100ms of the above channel model (see also Figure 10 for model details).

*Modeling the tactile evoked response*

Model simulations of tactile evoked responses were generated by a sequence of three inputs to SI, as in (1, 2). Based on this prior work, the first "feedforward", or bottom-up, input is simulated to arrive at pyramidal basal dendrites 25ms after stimulus onset, ostensibly from L4 by way of the sensory thalamus. Then, at 70ms, a top-down "feedback" input arrives from SII or higher-order thalamus. A subsequent basal input from L4 arrives 135ms poststimulus; however, we restricted our analysis of evoked responses to the first 140ms following the tactile stimulus because variability in the late input may depend on unmodeled interactions between SI and other cortical regions e.g. premotor cortex (20). See Supplemental Table 1 for parameters of evoked inputs.

We obtained substantially better agreement with evoked response curvature by smoothing the raw model evoked response with a 45ms Hamming window. A primary mismatch between model and data was the M70 depth, which is shallow in the model compared to the data (Figure 3C and 3D). Our primary conclusions hold true irrespective of this discrepancy.

*Model of beta event generation and hypothesized recruitment of long-time scale supragranular inhibition*

In a recent study, we showed that brief ~50ms bursts of excitatory synaptic input to supragranular layers from unspecified upstream regions can explain coarse spectrotemporal features of the beta event waveshape (15). Spiking in SI pyramidal cells is not needed to explain these features, which can arise due to the subthreshold nature of macroscale beta events, particularly those with sources in matrix or "modulatory" thalamus (21, 22). Our hypothesis that a primary source of burst inputs generating beta events is higher order (nonlemniscal) thalamus (15), a hypothesis bolstered by a recent study showing that nonlemniscal thalamic spikes generate subthreshold effects on both L2/3 and L5b pyramidal neurons in mice. These inputs also cause suprathreshold, basally-generated spikes in L54, concurrently recruiting VIP+ interneurons and potentially other 5HT3a+ interneurons in L2/3 (23).

The immediate action of nonlemniscal thalamus on cortex appears to be disinhibition and subthreshold excitation – a puzzle in light of our behavioral data. Yet, on longer timescales, strong excitation of rodent nonlemniscal homolog POm profoundly *reduces* sensory-evoked spiking in barrel cortex (24), suggesting that thalamus may act on cortex at multiple timescales by first facilitating and then suppressing activity. Although the cortical response to higher-order thalamic bursts has not been directly quantified, we hypothesize that the thalamic inputs responsible for beta events elicit long-timescale inhibition of cortical pyramidal neurons. Noting that the tactile suppression timescale corresponds to that of $GABA_{B1a}$ G-protein coupled inhibition, we furthermore propose that L2/3 neurogliaform cells (NGFCs) can mediate this effect.

One of the few cell classes known to act through $GABA_{B1a}$ receptors, NGFCs are represented among 5HT3a+ interneurons in L2/3 and are coupled to all other interneuron populations via gap junctions (25). Direct recruitment of NGFCs through nonlemniscal bursting is therefore plausible (23), but local neurogliaform cells can also be influenced to spike through electrical coupling -- synchronized VIP+ recruitment by a thalamic burst is arguably the single most likely means of activating an NGFC "circuit-breaker" in L2/3. These neurogliaform cells can act perisomatically on L2/3 pyramids, either synaptically (26, 27) or through volume transmission (25) and on middle-apical dendrites of L5 pyramidal cells (28).

Based on this proposed mechanistic framework, we simulate the inhibitory effects of $GABA_B$ in trials that include a prestimulus beta event, as follows. We drive the L2/3 inhibitory neurons strong enough to elicit a spike, and assume that in addition to impacting the L2/3 somata, there is an additional $GABA_B$ "synapse" from L2/3 inhibitory cells to the L5 pyramidal neuron middle-apical dendrite (see Figure 4B). This inhibition represents the L2/3 neurogliaform cell activation, as bulk GABA release is not explicitly modeled. These $GABA_B$ "synapses" onto the L5 apical dendrite were not present in simulations without a prestimulus beta event, but we did not alter somatic $GABA_B$ conductances between models with and without beta events (Figure 4). See Supplemental Table 1 for parameter values of the beta-generating drive, and for a comparison to earlier model parameters (29).

### Comparing MEG and Model Data

#### *Statistical analysis*

##### *Hypothesis testing*

Let $X$ be a data matrix with $X(i,t)$ representing the observation at time $t$ during trial $i$. Here $X(i,t)$ will be some property, such as current dipole or instantaneous phase, of a MEG signal from a single source-localized channel (real or modeled) at time $t$ on trial $i$. Let $L$ be vector of labels with $L(i)$ representing the label for trial $i$. Here $L(i) \in \{0,1\}$ will indicate a behavioral outcome or prestimulus event, such as a correct detection or a beta event, for trial $i$.

We use permutation tests to test the null hypothesis that the labels and the data at time $t$ are independent (or, more generally, that the labels are exchangeable given the data, at time $t$). For current dipole data our test statistic is the difference between the average current dipole on trials with label 1 and the average current dipole on trials with label 0. For instantaneous phase data our test statistic is the difference between the phase coherence on trials with label 1 and the phase coherence on trials with label 0.

Permutation tests work by randomly shuffling the trial labels and recomputing the test statistic for nonsensically labelled data many times in order to obtain a null distribution. The observed test statistic (on the correct trial labels) is compared to this null distribution in order to obtain a p-value. We report two-sided p-values in all cases and use at least 5000 permutations. See (30) for more details about hypothesis testing and permutation tests. We use the same trial label permutation for testing each time $t$ (i.e. the shuffling is performed on entire time-series, not independent time points). This is important for our method of controlling for multiple hypothesis tests, described next.

Because we are testing a separate null hypothesis at each time $t$, it is important to control for multiple hypothesis tests. We use a variant of the max-t method, which creates a global test statistic by taking the maximum (for the upper-tail) and minimum (for the lower tail) of the test-statistic over all times $t$, and then creates a null distribution (one for each tail) in the usual way by shuffling trial labels. The observed test statistic at each time $t$ is compared to these common, global null distributions in order to create adjusted p-values that provide strong control of the family-wise error rate. Rejecting those null hypotheses with adjusted p-values $\leq \alpha$ guarantees that the probability of zero false rejections is $\geq 1 - \alpha$. See (31, 32) for more details about permutation tests and multiple testing.

Our variant of the max-t method robustly standardizes the test statistics at each time $t$ prior to computing the maximum (or minimum) in order to more evenly distribute statistical power across all of the hypotheses. The specific details of our procedure can be found in (33) (We thank M. Harrison, Brown University, for sharing code for this procedure.)

### Supplementary Discussion

Beta Mediation of Learning

A natural pathway for beta events to induce learning is through the VIP+ interneuron system, known to be activated by higher-order thalamus (23) and recently shown to mediate NMDA-based long term potentiation (LTP) in pyramidal cells (34). Though not directly modeled, we hypothesized that the aforementioned VIP networks, when synchronized, also recruit neurogliaform cells via electrical synapses, which causes beta event suppression in our model. Interestingly, in rodent somatosensory L2/3 pyramidal cells, NMDA spike-mediated LTP, also with a thalamic source, had been found in the *absence* of somatic activity (35). This parallels our findings that L2/3 inhibition with a thalamic source results in a large dipole signal in the absence of somatic spiking (Figure 7).

Along these lines, it is interesting to note that increased somatic calcium flow has been reported under strong somatic inhibition during slow-wave sleep, where it was hypothesized to optimize learning (36). In that study, $GABA_A$-ergic parvalbumin positive interneurons were found responsible for the effect – however, $GABA_B$ activation tends to follow hyperactivation of $GABA_A$ interneurons as GABA overflows the synaptic cleft (37). Therefore, in addition to this VIP-NMDA mechanism, the collision between dendritic spikes and perisomatic suppression may lead to calcium transfer to the soma; such signaling cation fluxes are well-known to play fundamental and general roles in structural plasticity (38, 39). Electromotive forces generated in high voltage dendritic sodium spikes can presumably push calcium directly from the dendrite into the soma. Moreover, in the extreme case of long-duration dendritic bursts, the sheer voltage difference between compartments can also lead to an energy transfer large enough to cause perisomatic heating (see SI Appendix, Supplemental Calculation 1). Calcium influx due to momentary heating can then be effected through temperature-sensitive ion channels or through membrane capacitance changes, as evident in infrared stimulation protocols (e.g. 40).

Therefore, together with our modeling results, the literature appears to support two mechanisms — one short-term process via VIP - NMDA at the dendrite, and one long-term process via calcium at the soma — both of which might be linked by coupling through the Class 2 mechanism. It is possible that VIP-coupled NMDA may act as a short-term tag on a synaptic site, which could later be converted to permanent memory after spike destruction and subsequent somatic calcium influx, mediated by neurogliaform cells and/or soma-targeting $GABA_A$-ergic populations.

Modeling Assumptions, Limitations and Independence

This study builds from a body of prior MEG and modeling work, where we first showed that post-stimulus features of the tactile-evoked response in SI (i.e. the M70 amplitude and slope) alone could, in principle, account for correlates of tactile detection without considering prestimulus state (1). Later, we established that prestimulus low-frequency rhythms (i.e. the SI mu rhythm, comprised of 7-14 Hz alpha and 15-29 Hz beta rhythms) influence components of the evoked response through specific network mechanisms, including a strong inhibitory influence mediated by sensory evoked inhibition (2). However, in the latter study, we reported only on averaged data, and did not separate the effects of the alpha and beta components of the SI mu rhythm, nor did we investigate the relation of these effects to perception. Further studies showed alpha and beta have separable effects on perception and attention (18, 41). The current study is the first to look at circuit mechanism mediated perceptual effects specifically in the beta band.

The chosen SI model configuration is grounded in generalizable principles of cortical circuitry and known somatosensory cortical architecture (see Experimental Procedures). Some of the model assumptions create limitations in our conclusions, while many of the findings are independent of specific model choices.

One potential limitation is that we simulate only one type of $GABA_B$ receptor. We found that even at low densities, simulated $GABA_{B1a}$ channels in middle-apical dendrites of L5 pyramidal cells induced by beta-events can prevent these cells from firing during sensory stimulation. However, the primary target of L2/3 NGF activation on L5b apical dendrites is presumably the $GABA_{B1b}$ receptor, which inactivates calcium channels while admitting sodium spike propagation (28). As such, it is possible that L5 pyramidal spikes can be recruited by weak sensory stimuli in the absence of L2/3 recruitment when they are close to their firing threshold. Our model, and the assumed higher-order thalamic origin of burst events, also does not account for the higher-order thalamic recruitment of L5a pyramidal spikes observed in rodent slices (23).

Despite these assumptions, it is crucial to note that several of our main results, while dependent on our proposed beta-generating mechanism – namely, the ~50ms burst of subthreshold excitatory synaptic input to pyramidal neuron distal dendrites -- are essentially independent of free parameters in the model. First, the high-frequency coherence predicted during detected Class 1 responses relies only on the ~5ms L2/3 doublet interval observed in our model, which has been noted in awake rodents and may possibly have a correlate in primates (42, 43). Second, non-detected Class 2 waveforms should occur in *any* model that contains dendritic geometry, dendritic spikes, and strong perisomatic inhibition. Third, while we've assumed beta events are mediated by higher-order thalamus, it is possible that top-down corticocortical connections play a role in generating beta events. The functional distinction of the source of the distal drive does not change the fundamental findings of our study, which identifies neural circuit mechanisms generating beta-mediated evoked response correlates of perception within a canonical cortical unit.
